# Therapeutic targeting of differentiation state-dependent metabolic vulnerabilities in DIPG

**DOI:** 10.1101/2022.03.01.482555

**Authors:** Nneka E. Mbah, Amy L. Myers, Chan Chung, Joyce K. Thompson, Hanna S. Hong, Peter Sajjakulnukit, Zeribe C. Nwosu, Mengrou Shan, Stefan R. Sweha, Daniella D. Maydan, Brandon Chen, Li Zhang, Brian Magnuson, Zirui Zui, Daniel R. Wahl, Luigi Franchi, Sameer Agnihotri, Carl J. Koschmann, Sriram Venneti, Costas A. Lyssiotis

## Abstract

H3K27M diffuse intrinsic pontine gliomas (DIPG) exhibit cellular heterogeneity comprising less-differentiated, stem-like glioma cells that resemble oligodendrocyte precursors (OPC) and more differentiated astrocyte (AC)-like cells. H3K27M DIPG stem-like cells exhibit tumor-seeding capabilities *in vivo*, a feature lost or greatly diminished in the more differentiated AC-like cells. In this study, we established isogenic *in vitro* models of DIPG that closely recapitulated the OPC-like and AC-like phenotypes of DIPG cells. Using these tools, we performed transcriptomics, metabolomics, and bioenergetic profiling to identify metabolic programs operative in the different cellular states. From this, we defined new strategies to selectively target metabolic vulnerabilities within the specific tumor populations. Namely, we showed that the AC-like cells exhibited a more mesenchymal phenotype and were thus sensitized to ferroptotic cell death. In contrast, OPC-like cells upregulated cholesterol metabolism and mitochondrial oxidative phosphorylation (OXPHOS) and were accordingly more sensitive to statins and OXPHOS inhibitors. Additionally, statins and OXPHOS inhibitors showed efficacy and extended survival in preclinical orthotopic models established with stem-like H3K27M DIPG cells. Together, this study demonstrates that cellular subtypes within DIPGs harbor distinct metabolic vulnerabilities that can be uniquely and selectively targeted for therapeutic gain.

## INTRODUCTION

Diffuse intrinsic pontine gliomas (DIPG) are treatment-resistant and uniformly fatal pediatric brain tumors. The prognosis of this brainstem tumor is dismal with a median overall survival of 9-12 months from diagnosis^1, 2^. Radiotherapy is the only therapy that has proven benefit in this patient population. Several clinical trials with chemotherapy have failed to demonstrate any additional survival benefit over radiation alone^3, 4^. It is therefore imperative to identify novel strategies for targeting this aggressive and devastating disease.

Recent advances delineating the molecular underpinnings of DIPG revealed that approximately 80% of DIPGs and diffuse midline gliomas (DMGs) harbor a recurrent somatic lysine-to-methionine mutation at position 27 of histone H3.1 (*HIST1H3B*) or H3.3 (*H3F3A*), collectively called H3K27M. H3K27M results in global hypo-methylation on H3K27 sites, epigenetic dysregulation, and altered gene transcription^1, 5–8^. Other activating mutations and aberrant gene expression patterns identified in DIPG include *ACVR1, TP53,* and *ATRX* mutations, and overexpression of transcriptional factors OLIG1 and OLIG2 ^9–11^.

Single-cell RNA sequencing studies analyzing over 3,000 H3K27M cells from six primary H3K27M gliomas found that H3K27M DIPGs hijack developmental programs regulating lineage differentiation of neural stem cells. Consequently, tumor cells undergo developmental arrest and are locked in a less-differentiated state^12^. This study also revealed that DIPGs contain a heterogeneous population of cells, where the majority of tumor cells harbor characteristic markers of ‘oligodendrocyte precursor cells’ (OPC-like), with stem-like and higher renewal potential. The more differentiated-like H3K27M tumor cells exhibit an astrocytic-like (AC-like) phenotype and represent a minority^12^. Of note, in the context of DIPG cellular heterogeneity, the H3K27M OPC-like cells are hypothesized to be the putative drivers of tumor growth and aggressiveness and possess *in vivo* tumor-initiating potential compared to more differentiated cells^12–15^.

Metabolic reprogramming is a hallmark of cancer that influences every aspect of cancer biology^16, 17^. Indeed, in comparison to H3 wild-type (H3WT) tumor cells, H3K27M DIPG cells enhance metabolic programs including glycolysis, glutaminolysis, and the tricarboxylic acid (TCA) cycle, as well as increase the generation of α-ketoglutarate (α-KG). Of note, enhanced α-KG production in H3K27M DIPG was shown to be critical for the maintenance of the preferred global H3K27 hypomethylation status, indicating the important interplay between metabolic reprogramming and epigenetic regulation^18^. Despite these emerging studies of metabolism in H3K27M DIPG, the role of dysregulated metabolism, particularly in the context of how metabolism impacts stemness and tumorigenicity in H3K27M DIPG, remains largely unknown. Indeed, differential metabolic reprogramming can potentially regulate cancer stemness, differentiation, and cell fate ^19^. As a result, many aspects of rewired metabolism can provide novel therapeutic liabilities within tumors that can be effectively leveraged for therapy.

In this study we sought to elucidate the metabolic dependencies operative in both the stem-like tumorigenic H3K27M gliomas and the more differentiated cell state. By applying a systems biology approach that incorporated metabolomics, transcriptomics, and biochemical analyses, we uncovered several nodes of dysregulated metabolic and signaling pathways in the tumorigenic OPC-like versus (vs.) the more differentiated AC-like DIPG populations. This study collectively illustrates that DIPGs harbor perturbations in metabolic programs that can be exploited for therapeutic benefits.

## RESULTS

### *In vitro* modeling of the differentiation state of H3K27M midline gliomas

To study metabolic vulnerabilities associated with the distinctly heterogenous H3K27M DIPG, we generated isogenic gliomaspheres (GS) and differentiated glioma cells (DGC) from three patient-derived H3K27M DIPG cell lines: DIPG-007, DIPG-XIII and SF7761. It has been previously established that the DIPG GS culture conditions (serum-free) enrich for malignant cells that are less-differentiated and stem-like^20, 21^. Moreover, DIPG GS readily establish tumors following stereotactic injection into the pons of mice^12^. In contrast, culturing the tumor cells in the presence of serum induces differentiation of H3K27M malignant glioma cells (i.e., DGC) and an associated loss or substantially diminished *in vivo* tumorigenicity^12^. Furthermore, in comparison to the DGC, the GS model most closely recapitulates the phenotype of DIPG tumor-xenografts and primary patient tumors^12, 20–22^.

We therefore maintained the isogenic cell lines either as tumorigenic GS cultures (i.e., unattached 3-D spheres cultured in serum-free media containing B27 supplements and growth factors) or differentiated malignant cells (i.e., adherent monolayers cultured in the presence of serum) (**Figure 1A; Supplementary Figure 1A**). Evaluation of the growth kinetics of GS vs. DGC revealed that DIPG-007 and SF7661 GS proliferated at a markedly higher rate than their DGC counterparts, indicating that the less differentiated and stem-like GS populations are far more proliferative. The differentiation state did not influence proliferation rates in the DIPG-XIII cells (**Figure 1B**).

**Figure 1:**
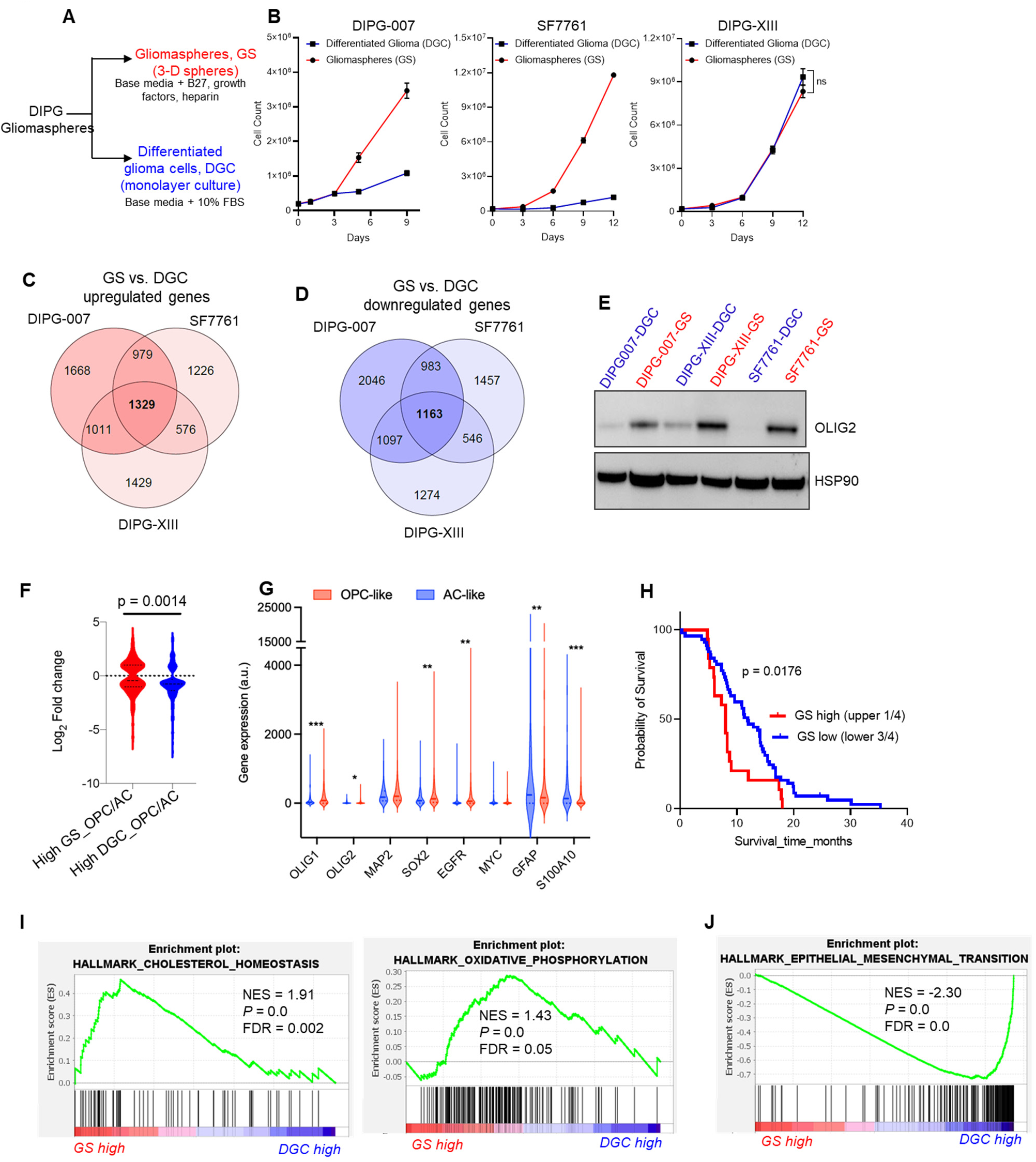
*In vitro* models of H3K27M DIPG molecularly mimic *OPC-like* and *AC-like* DIPG and exhibit distinct gene expression programs. **A)** Schematic depicting the generation of isogenic DIPG gliomaspheres (GS) and differentiated glioma cells (DGC): GS (3-D floating spheres) were cultured in serum-free tumor stem cell (TSM) media containing growth factors and supplements. Monolayer (adherent) DGCs were generated by dissociating GS to single cells and culturing for up to 14 days in TSM media containing 10% fetal bovine serum (FBS). **B)** Growth kinetics of DIPG-007, SF7761 and DIPG-XIII GS vs. DGC. **C-D)** Venn diagrams illustrating the overlap of significantly upregulated **(C)** and downregulated **(D)** genes (adjusted p-value <0.0001) in GS vs. DGC across three isogenic lines. **E)** Western blot for Olig2 in GS and DGC isogenic pairs of patient-derived DIPG cell lines. HSP90 was used as a loading control. **F)** Analysis of the gene expression signature of DIPG GS vs. DGC cross-referenced with patient gene signature capturing OPC-like, AC-like gene signatures. **G)** Differential expression of DIPG genes in OPC-like and AC-like tumors from patient single-cell RNAseq. **H)** Survival analysis of DIPG/DMG patients based on “GS high” versus “GS low” gene signature. **I-J)** Representative “Hallmark” gene set enrichment analysis (GSEA) indicating pathways that are **I)** upregulated and **J)** downregulated in isogenic DIPG GS vs. DGC. GSEA plots show enrichment scores and include values for normalized enrichment score (NES), nominal p-value (*P*), and false discovery rate (FDR) q-value.

### Transcriptomic analyses revealed GS are OPC-like and DGC are AC-like

To characterize the molecular features of the DIPG GS and DGC, we performed RNA-sequencing (RNA-seq) to compare the gene expression patterns of GS vs. DGC in these three isogenic cell lines. Principal component (PC) analysis of the RNA-seq data revealed that the cell lines clustered based on cell line differences and cellular differentiation status (namely, DGC or GS) (**Supplementary Figures 1B-C)**. PC1 vs. PC2 was shown to be driven by original cell line differences and clustered based on cell line, irrespective of the differentiation state (**Supplementary Figure 1B**). PC3 distinguished between GS and DGC, wherein the isogenic lines clustered separately, suggesting that PC3 was influenced by the cellular differentiation status. Strikingly, PC3 revealed that GS vs DGC gene expression patterns were more strongly separated in DIPG-007 and SF7761 than in DIPG-XIII (**Supplementary Figure 1C**). These observations suggest a potentially marked difference in the resultant phenotype of GS vs. DGC in DIPG-007 and SF7761, which is less pronounced in DIPG-XIII.

We analyzed genes that were significantly altered (adjusted p-value < 0.0001) in GS vs. DGC across the three cell lines (DIPG-007, 10276 total genes; SF7761, 9242 total genes; and DIPG-XIII, 8425 total genes) and found 1329 genes and 1163 genes to be commonly upregulated and downregulated, respectively (**Figures 1C-D**). The top 50 consistently upregulated and downregulated genes common to the three isogenic lines revealed distinct gene signatures associated with GS vs. DGC tumor populations. (**Supplementary Figure 1D-E**). Next, we examined the gene expression of individual DIPG stemness and differentiation markers. GS lines upregulated oligodendrocyte transcription factor 2 (*OLIG2*) (**Supplementary Figure 2A**), the gene that encodes a transcription factor typically overexpressed in DIPG and critical for the establishment of DIPG tumors *in vivo*^24^. This difference was also observed at the protein level (**Figure 1E**). In addition, other DIPG markers like oligodendrocyte transcription factor 1 (*OLIG1)* and microtubule associated protein 2 (*MAP2)* were higher in GS compared to GDC (**Supplementary Figures 2B-C**). SRY-box transcription factor 2 (*SOX2)*, epidermal growth factor receptor (*EGFR)*, and Myc proto-oncogene (*MYC)* are genes whose upregulation is associated with glioma stemness. Remarkably, these genes showed consistently higher expression in DIPG-007 and SF7761 GS compared to their DGC counterparts, and this contrasted with the modestly lower expression in DIPG-XIII GS vs. DGC (**Supplementary Figures 2D-F**). Analysis of glial fibrillary acidic protein (*GFAP*), vimentin (*VIM*), and S100 calcium binding protein A10 (*S100A10*), which are typically associated with astrocytic-like differentiated DIPG^25–27^, revealed substantial upregulation in DGC compared to GS across the three lines (**Supplementary Figures 2G-I**).

Malignant H3K27M gliomas have been reported to include tumor cell types that exhibit four gene signatures: namely, i) OPC-like cells, ii) cell cycle (CC), iii) oligodendrocytes (OL), and iv) AC-like cells^12^. We cross referenced our bulk RNA-seq gene expression data with that of the published single-cell RNA-seq dataset^12^ (GSE102130) that described the four gene signatures. We found that the gene expression pattern observed in the GS showed increased enrichment for the OPC-like gene signature, while the gene signature of the DGCs was consistent with an AC-like phenotype (**Figure 1F**). Interestingly, individual gene expression trends observed in our *in vitro*-generated isogenic GS vs. DGC cells (**Supplementary Figures 2A-I**) were also recapitulated in the OPC-like vs. AC-like H3K27M cells defined in this published single-cell RNA-seq dataset (**Figure 1G)**.

Collectively, these data suggest that *in vitro* generated GS largely represent the OPC-like DIPG phenotype, which is known to be less-differentiated, stem-like, and exhibit tumor-propagating potential *in vivo*, while DGC represent the more differentiated AC-like phenotype. Accordingly, the GS and DGC *in vitro* models developed herein molecularly mimic two distinct and predominant populations in the heterogenous H3K27M DIPG tumor.

### GS gene signature predicts decreased survival of DIPG/DMG patients

To determine the clinical relevance of the GS gene signature in predicting disease outcome and survival of patients with H3K27M DIPG, we mined a patient dataset from Mackay et al.^28^ containing gene expression and survival data for 76 H3K27M DIPG and DMG patients. We segregated patients into “high GS” vs. “low GS” gene expression categories using unbiased K-means clustering and applied Kaplan-Meier survival analysis to define upper quartile as “GS high” vs. “GS low” tumors. The results revealed that patients with “GS high” tumors showed a significantly decreased survival in comparison to patients in the “GS low” category within H3K27M tumors (**Figure 1H**). This result supports our observation that “GS” gene signature, which recapitulates those of the less differentiated OPC-like cells, represents the more aggressive and tumorigenic cell-state of DIPG.

### OPC-like GS upregulate cholesterol metabolism and oxidative phosphorylation

To interrogate the gene expression programs that distinguish the cell state among our isogenic pairs, we performed gene set enrichment analyses (GSEA) on each of our RNA-seq dataset from the three isogenic lines. Across the three lines, DIPG GS vs. DGC upregulated genes were associated with the MYC pathway, PI3K/MTORC1 signaling, G2M checkpoint, DNA repair, and E2F signaling (**Supplementary Figure 3A**). These pathways have been previously reported to be upregulated in primary patient DIPG tumors and xenografts^28–30^, thereby providing further confidence in our DIPG models and analyses. Furthermore, in at least two of the three isogenic lines, we observed considerable DIPG GS enrichment of metabolic pathways, namely cholesterol homeostasis and mitochondrial oxidative phosphorylation (OXPHOS) (**Figure 1I**). In contrast, DIPG DGC upregulated genes associated with epithelial-mesenchymal transition (EMT) (**Figure 1J**), xenobiotic metabolism, inflammatory response, and transforming growth factor (TGF)β signaling (**Supplementary Figures 3B**). These findings suggested that tumorigenic OPC-like GS may exhibit enhanced reliance on cholesterol metabolism and mitochondrial OXPHOS programs, which could represent an actionable metabolic vulnerability.

### Steady-state metabolomics reveal distinct metabolic profiles of GS vs. DGC

Given the metabolic signatures evident in the transcriptomic profiling, we performed liquid chromatography-coupled mass spectrometry (LC/MS)-based metabolomics^31^ to gain a deeper understanding of metabolic differences between the DIPG cellular differentiation states. These data revealed that GS and DGC exhibit distinct metabolic landscapes (**Supplementary Figure 4**). By taking the average of the three isogenic lines, we found that compared to DGC, GS showed a significant difference in nucleotide metabolism, lipid and sterol biosynthesis, and amino acid metabolism (**Figures 2A-B**). Further, among the ∼223 metabolites measured, we found 70 that were significantly altered between GS and DGC. Of these, 45 metabolites were highly increased, and 25 metabolites were decreased in GS compared to DGC (**Figure 2C**).

**Figure 2:**
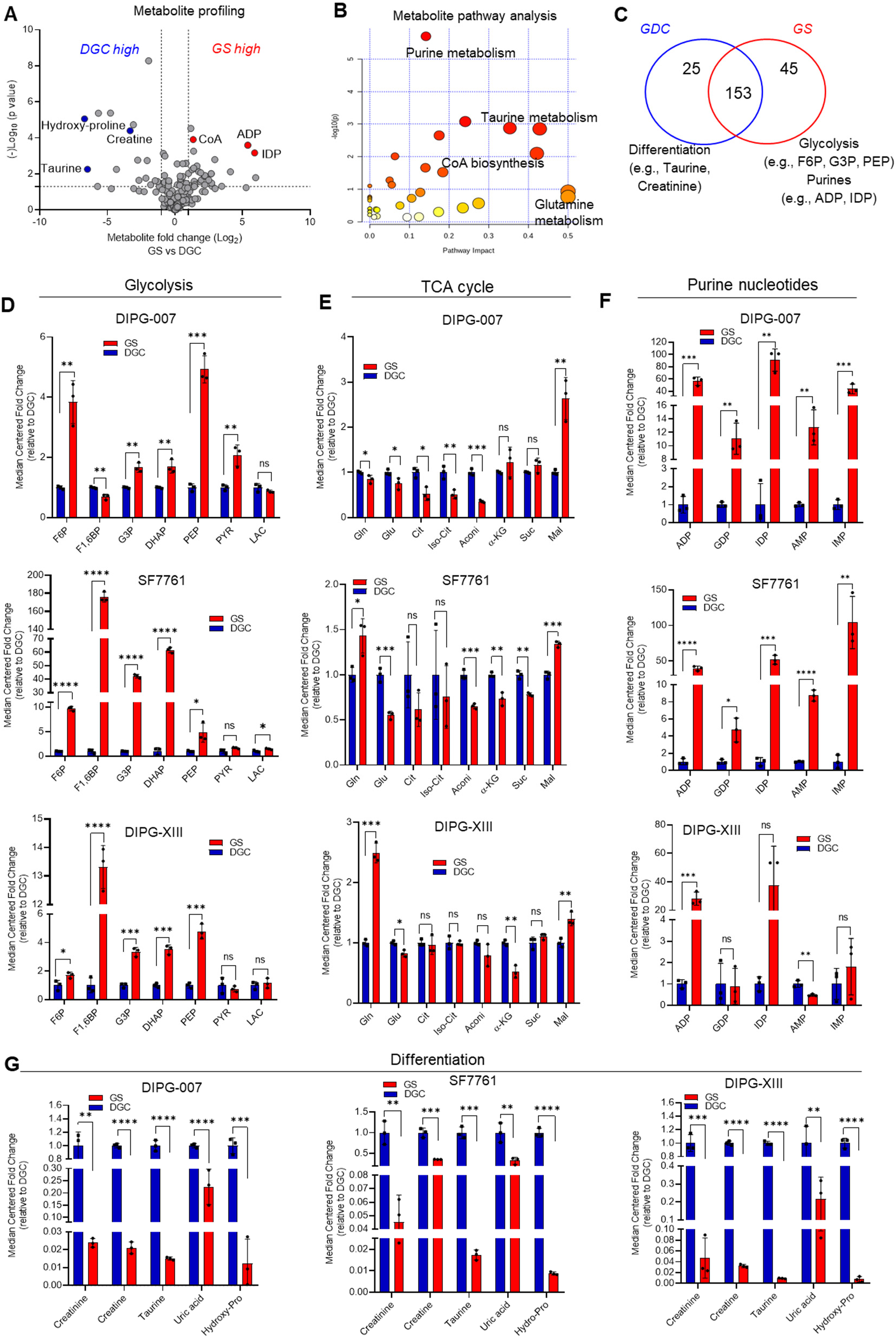
Metabolomic profiling of isogenic GS and DGC gliomas reveal differentiation state-dependent metabolic features. **A)** Volcano plot indicating differential metabolite profiles in GS vs. GDC, presented as the average expression value from the three cell lines; DIPG-007, SF7761, and DIPG-XIII. **B)** Metabolic pathway enrichment as determined using MetaboAnalyst based on differential metabolite abundance in GS vs. DGC. **C)** Venn diagram indicating the number of metabolites significantly altered in GS vs. DGC across all three cell lines. **D-F)** Differential abundance of select metabolites for **D)** glycolysis, **E)** tricarboxylic acid (TCA) cycle, and **F)** purine nucleotides in DIPG-007, SF7761, and DIPG-XIII GS versus DGC. **G)** Highly enriched metabolites in DIPG DGC vs. GS. Metabolite levels expressed as median centered fold change of GS relative to DGC across the three isogenic lines (ns = not significant; * p < 0.05; ** p < 0.01; *** p < 0.001; **** p < 0.0001). Error bars represent mean ± SD. Source data are provided as a Supplementary Table 1. F6P, fructose-6-phosphate; F1,6BP, fructose-1-6-bisphosphate; G3P, glycerol-3-phosphate; DHAP, dihydroxyacetone phosphate; PEP, phosphoenolpyruvate, PYR, pyruvate; LAC, lactate; Gln glutamine; Glu, glutamate; Cit, citrate; Iso-cit, Isocitrate; Aconi, cis-Aconitate; αKG, alpha-ketoglutarate; Suc, succinate; Mal, malate; ADP, adenosine diphosphate; GDP, guanosine diphosphate; AMP, adenosine monophosphate; IMP, inosine monophosphate; IDP, inosine diphosphate; Hydro-Pro, hydroxy-proline.

Several metabolites were significantly altered in glycolysis, the TCA cycle, and purine biosynthesis pathway (**Supplementary Figure 5A**). Glycolytic metabolites were generally more abundant in GS compared to DGC (**Figure 2D**), with several differences being greater than 10- fold, particularly those in the preparatory phase of glycolysis. Pyruvate and lactate, products of aerobic and anaerobic glycolysis, respectively, showed either no significant difference between the isogenic states or were modestly altered (**Figure 2D**). Glycolysis connects to the TCA cycle via the generation of acetyl CoA from CoA and pyruvate. Metabolites in the TCA cycle were generally decreased in DIPG-007 and SF7761 GS compared to GDC, while few significant differences were observed in the DIPG-XIII isogenic pair. An exception was malate, which was significantly increased in GS across the three isogenic lines (**Figure 2E**). The metabolomics studies also revealed increased levels of CoA and carnitine, key metabolites and rate-limiting substrates in lipid and sterol biosynthetic pathways (**Supplementary Figure 5B**).

### Purine nucleotides are enriched in OPC-like GS

Of the phosphorylated purine species detected, markedly higher levels of purine nucleotide pools (> 50-fold in several cases) were observed in GS compared to DGC in DIPG-007 and SF7761, including adenosine monophosphate (AMP), guanosine monophosphate (GMP), inosine monophosphate (IMP), inosine diphosphate (IDP) and adenosine diphosphate (ADP) (**Figure 2F**). With the exception of ADP, such differences were not observed in DIPG-XIII (**Figure 2F**). Increased expression of genes encoding purine pathway enzymes were similarly observed (**Supplementary Figure 5C**), including phosphoribosyl pyrophosphate synthetase 2 (*PRPS2)* which converts ribose-5-phosphate to phosphoribosyl pyrophosphate; adenylosuccinate synthase 2 (*ADSS*) which converts IMP to adenylosuccinate; adenylosuccinate lyase (*ADSL*) which converts adenylosuccinate to AMP; inosine monophosphate dehydrogenase 1 (*IMPDH1*) which converts IMP to xanthine monophosphate (XMP); and guanine monophosphate synthase (*GMPS*) which converts XMP to GMP. Indeed, increased purine nucleotides pools have been demonstrated to be an intrinsic characteristic of brain tumor-initiating glioma cells^32^.

### AC-like DGC accumulate metabolites associated with cellular differentiation

Comparison of metabolite abundance in DGC relative to GS revealed increases on the order of 10-fold for taurine, creatine, creatinine, uric acid, and hydroxy proline (**Figure 2G**). These data were provocative because taurine, creatine, and creatinine have been shown to be elevated in oligodendrocytes generated by inducing differentiation of primary OPCs using triiodothyronine (T3)^33^. Moreover, exogenous taurine was shown to promote drug-induced differentiation of primary OPC cells to OLs, and is presumed to be synthesized by cells to promote lineage differentiation^33^.

### AC-like DGC upregulate EMT pathway genes and are vulnerable to ferroptosis

The results from our metabolomics studies identified several nodes of metabolism that differ between GS and DGC. We next sought to assess if these differences in metabolic programming provide therapeutic vulnerabilities. GSEA analysis illustrated that DGC upregulated genes associated with epithelial mesenchymal transition (EMT) and TGFβ signaling (**Fig 1J; Supplementary Figure 3B**). This EMT signature was characterized by a general increase in expression of EMT marker genes, including transforming growth factor beta 2 (*TGFB2),* vascular cell adhesion molecule 1 (*VCAM1*), snail family transcriptional repressor 1 (*SNAI1*), matrix metallopeptidase 2 (*MMP2*) and matrix metallopeptidase 11 (*MMP11*) (**Figure 3A**). Several recent studies have illustrated that the mesenchymal state of cancer cells exposes a vulnerability to ferroptosis - a form of metabolic-stress cell death induced by inhibiting the GPX4 lipid peroxidase pathway^34, 35^. Ferroptosis can be induced by genetic or pharmacological manipulations that impair cystine uptake, block glutathione (GSH) synthesis, or directly inhibit activity of the central lipid peroxidase, GPX4^36^ (**Figure 3B**). Based on this knowledge, we hypothesized that DGC, owing to its high mesenchymal gene signature, would be susceptible to GPX4 inhibition and ferroptosis.

**Figure 3:**
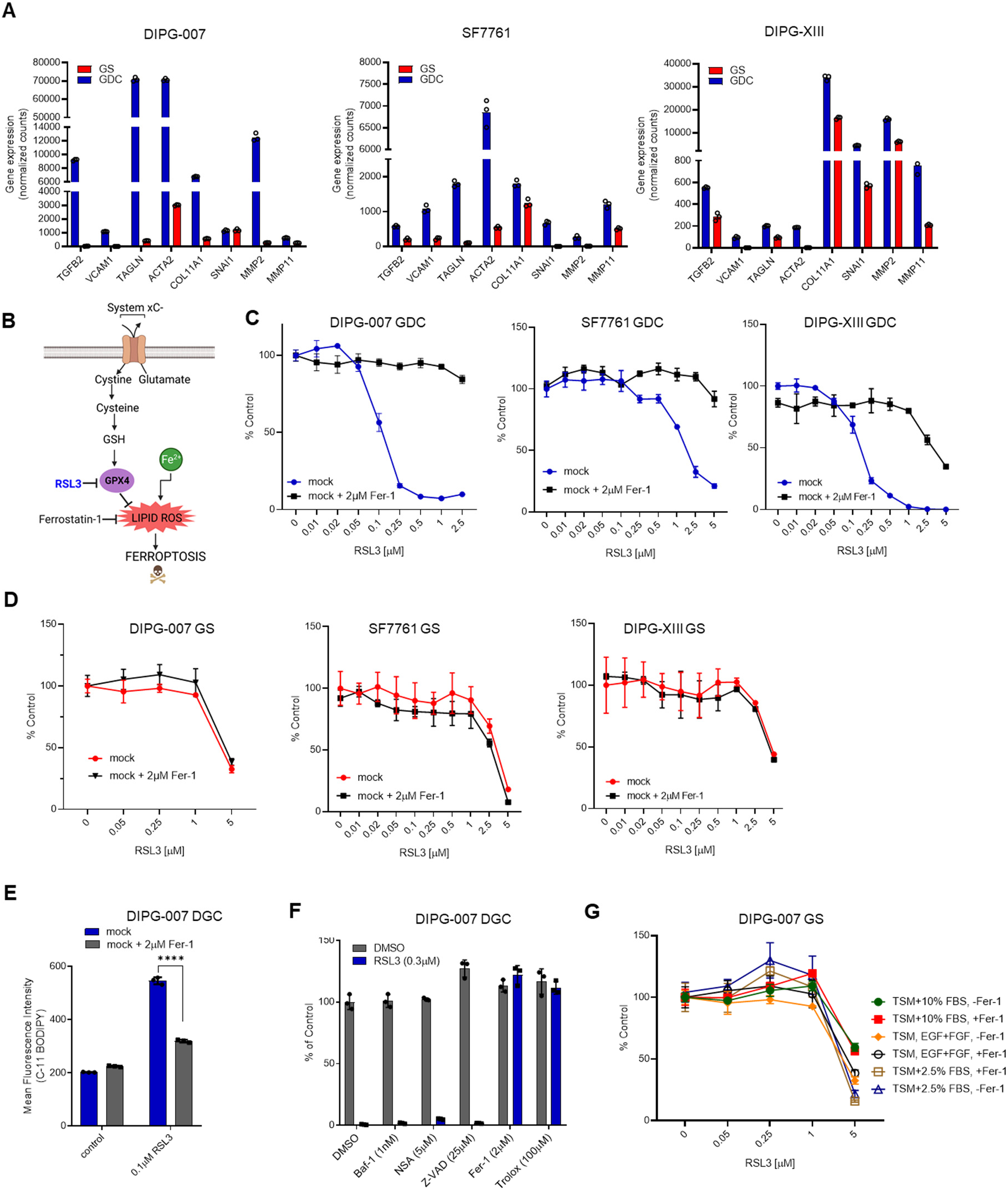
Ferroptosis is a metabolic vulnerability of AC-like DGC. **A)** Differential expression of genes associated with epithelial to mesenchymal transition (EMT) in GS vs. DGC across DIPG-007, SF7761, and DIPG-XIII models. **B)** Simplified scheme depicting the role of GPX4 in ferroptosis. **C, D)** Dose-response of DIPG-007, SF7761, and DIPG-XIII **C)** DGC, and **D)** GS treated with vehicle (0.1% DMSO) or RSL3 with or without 1h pre-treatment with 2µM ferrostatin-1 (Fer-1). Cell viability was assessed 48 hours post drug treatment via Cell Titer Glo 2.0. **E)** Flow cytometry assessment of lipid ROS in DIPG-007 DGC treated with either vehicle (DMSO 0.1%) or 1µM RSL3 for 6 hours, with or without 1h pretreatment with 2µM Fer-1. C-11 BODIPY dye used to quantify intracellular lipid ROS and data expressed as mean ± SD of mean fluorescent intensity (MFI) (****p < 0.0001). **F)** Cell viability of DIPG-007 DGC cultured in vehicle (0.1% DMSO) or 1µM RSL3 in the presence of 2μM Fer-1, 100μM Trolox, 2.5μM Necrosulfonamide (NSA), 25μM ZVAD-FMK (Z-VAD), or 1nM bafilomycin A1 (Baf-1). Viability was assessed 48 hours post drug treatment via Cell Titer Glo 2.0. Data expressed as a percentage of control (vehicle). Error bars represent mean ± SD. **G)** Cell viability of freshly dissociated DIPG-007 GS cultured in either GS growth media (TSM + growth factors EGF and FGF), DGC growth media (TSM + 10% FBS), or GDC reduced serum media (TSM + 2.5% FBS), followed by treatment with either vehicle (DMSO 0.1%) or different doses of RSL3 with or without 1h pre-treatment with 2µM Fer-1. Cell viability was assessed 48 hours post drug treatment via Cell Titer Glo 2.0. Data expressed as a percentage of control (vehicle). Error bars represent mean ± SD.

Treatment of DIPG-007, SF7761, and DIPG-XIII DGC with the GPX4 inhibitor RSL3 led to profound cell death by 48 hours (**Figure 3C**). To determine whether the RSL3-induced cell death was indeed ferroptotic in nature, we demonstrated that pre-treating cells with the lipophilic antioxidant ferrostatin-1 (Fer-1; a well-established inhibitor of ferroptosis^37^) rescued cell death.

In contrast to DGC, GS cells underwent RSL3-induced cytotoxicity at much higher concentrations, and more importantly, this effect could not be rescued by Fer-1. The lack of rescue illustrates a ferroptosis-independent mechanism of cell death (**Figure 3D**). Next, we assessed if RSL3 could induce lipid oxidation, a classic hallmark of ferroptosis, in DGC. Indeed, treatment of DIPG-007 with RSL3 resulted in increased accumulation of lipid reactive oxygen species (lipid ROS), as measured by C-11 BODIPY, which could be mitigated by pre-treating cells with Fer-1 (**Figure 3E**).

To rule out other avenues of RSL3-induced cytotoxic cell death, DGC were pre-treated with antioxidants and ferroptosis inhibitors (trolox, Fer-1), z-vad-fmk (apoptosis inhibitor), bafilomycin-A1 (autophagic cell death inhibitor), or necrosulfonamide (necroptosis inhibitor). Only the antioxidants rescued cell death induced by RSL3 in DIPG-007 DGC (**Figure 3F**), indicating a ferroptotic-specific mechanism of cell death.

As an important control, we also investigated whether the resistance of GS to ferroptosis was the result of culture media composition. To test this, we cultured freshly dissociated GS in serum-containing DGC media and assessed RSL3-induced ferroptosis. GS displayed a similar level of resistance to ferroptosis regardless of the media formulation (**Figure 3G**). These results illustrate that the growth medium itself does not directly impact the susceptibility of DIPG cells to ferroptosis. Rather, our data suggest that AC-like DGC have undergone the process of differentiation and, thereby, harbor cell-intrinsic qualities that promote sensitivity to ferroptosis. This is consistent with our hypothesis that the mesenchymal state of DGC sensitizes them to GPX4 inhibition-induced ferroptosis.

### Cholesterol biosynthesis and mitochondrial OXPHOS are metabolic vulnerabilities in OPC-like GS

Cholesterol homeostasis and mitochondrial OXPHOS were the top upregulated metabolic pathways by gene expression analysis in OPC-like GS (**Figure 1I**). Accordingly, we investigated the sensitivity of GS vs. DGC to OXPHOS inhibitors (Phenformin, Metformin, IACS-010759) or cholesterol biosynthesis inhibitors (statins). *In vitro* cultured DIPG-007 GS and DGC treated with increasing drug concentrations revealed that GS were strikingly and selectively more sensitive to Metformin, Phenformin, and IACS-010759 (**Figures 4A-C**), as well as to statins (Atorvastatin, Fluvastatin, and Pitavastatin) (**Figures 4D-F**). Of the five clinically available lipophilic statins, Pitavastatin, was most potent at reducing viability of DIPG-007 *in vitro* (**Supplementary Figure 6L**). Similar observations were made in SF7761 cells (**Suppl Figures 6A-D**). Neither DIPG-XIII nor DIPG-XIII-P* (a subclone of the DIPG-XIII that exhibits increased aggressive growth in vivo) demonstrated a consistent differential response to OXPHOS inhibitors. In contrast, their differential response to statins ranged from modest (DIPG-XIII) to substantial (DIPG-XIII-P*) (**Supplementary Figures 6E-H**). Notably, the murine cell line of <u>P</u>53 <u>P</u>DGFRA H3<u>K</u>27M DIPG (PPK cells), which has been demonstrated to replicate the H3K27M DIPG biology ^38^ and was similarly established in culture as either GS or DGC, also displayed differential sensitivity to Phenformin and Pitavastatin (**Supplementary Figures 6J-K**).

**Figure 4:**
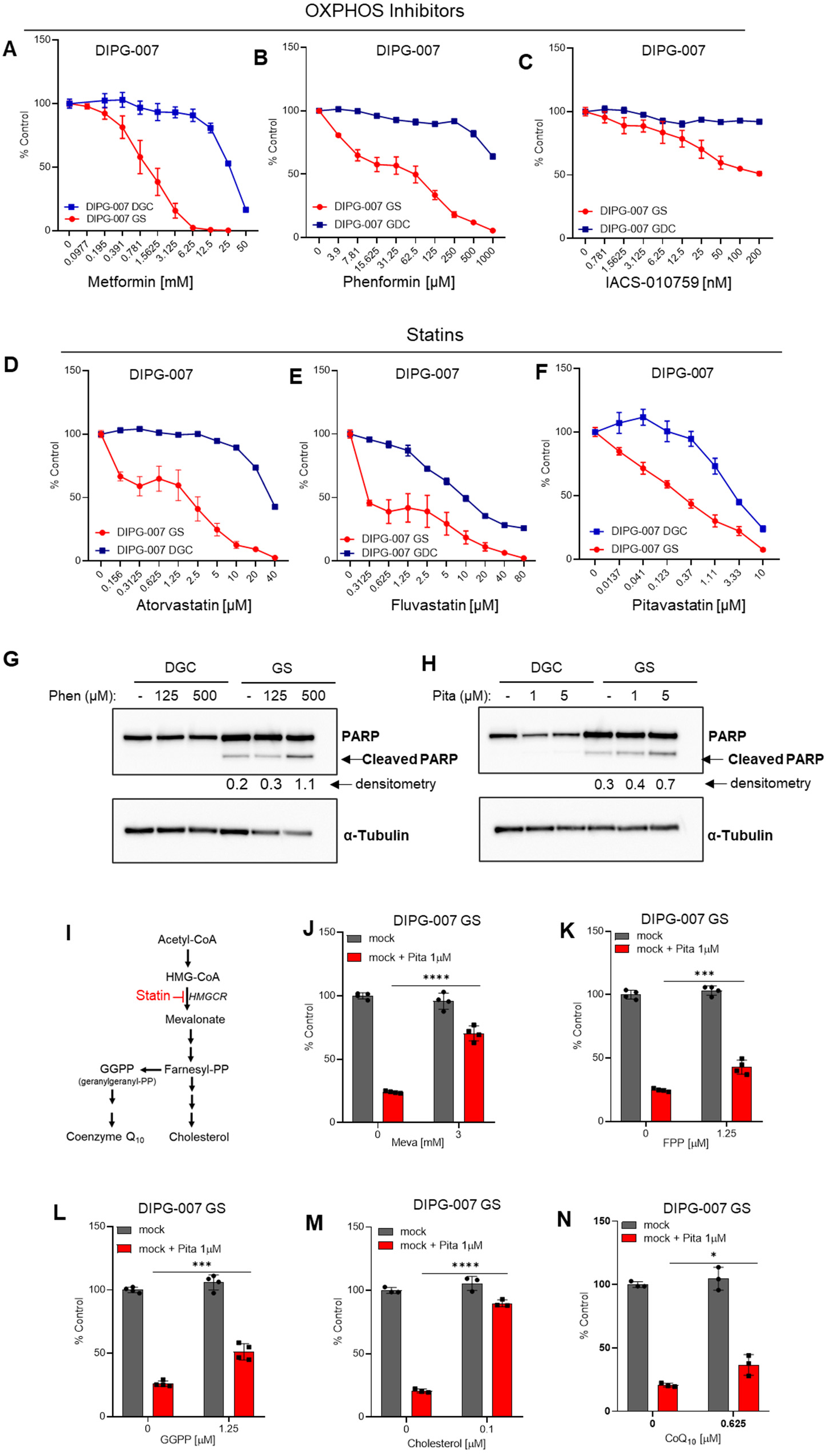
Cholesterol homeostasis and OXPHOS are targetable vulnerabilities in OPC-like GS. **A-F)** Dose-response curves for DIPG-007 GS vs. DGC treated with indicated concentrations of mitochondrial OXPHOS and complex I inhibitors **A)** Metformin, **B)** Phenformin, and **C)** IACS-010759 (IACS), or statins **D)** Atorvastatin, **E)** Fluvastatin, and **F)** Pitavastatin for 3 days. Cell viability assayed using the Cell Titer Glo 2.0 reagent and results expressed as a percent of control (vehicle, 0.1% DMSO) and mean ± SEM. **G, H)** Western blot of PARP cleavage (apoptosis indicator) and Tubulin (loading control) in **G)** Phenformin treated and **H)** Pitavastatin treated-DIPG007 DGC and GS. Cells were treated for 2 days at the indicated concentration. **I)** Schematic of sterol biosynthesis pathway indicating key intermediates in the biosynthesis of cholesterol and coenzyme Q_10._ **J-N)** Cell viability of DIPG-007 GS following treatment with vehicle (0.4% DMSO) or Pitavastatin (Pita) with and without co-treatment with **J)** Mevalonate (Meva), **K)** farnesyl pyrophosphate (FPP), **L)** geranylgeranyl pyrophosphate (GGPP), **M)** Cholesterol (Chol), and **N)** Coenzyme Q_10_ (CoQ_10_) at the indicated concentrations. Cell viability assayed at 3 days post-treatment using the Cell Titer Glo 2.0 reagent and results expressed as a percent of control and mean ± SD (ns = not significant; * p < 0.05; ** p < 0.01; *** p < 0.001; **** p < 0.0001).

To determine the mechanism of cytotoxicity induced by OXPHOS inhibition (Phenformin) and statins (Pitavastatin), we treated DIPG-007 DGC and GS with equal concentrations of these compounds and assessed PARP cleavage via western blot as a readout for apoptotic cell death. The results showed modestly elevated levels of PARP cleavage in DIPG-007 GS but not in DGC, demonstrating that the cytotoxic effects of statins and OXPHOS inhibition are at least partially the result of induction of apoptosis cell death (**Figures 4G-H**).

In an attempt to invoke an even more potent cytotoxic effect using both pathway inhibitors, DIPG-007 GS and murine PPK GS were treated with increasing doses of single agent Pitavastatin, Phenformin, or the combination with the IC_25_ or IC_50_ of the respective combinatorial compound. The results revealed modest sub-additive activity in DIPG-007 (**Supplementary Figures 7A-B**). No additional cytotoxic benefit was found in murine PPK GS treated with either Metformin or Phenformin in combination with Pitavastatin (**Supplementary Figures 7C-7F**).

### OPC-like GS depend on the sterol biosynthesis pathway for cholesterol

Based on the upregulation of cholesterol metabolism gene expression (**Figure 1I**, **Supplementary Figures 8A-C**) and the robust sensitivity of GS to statins, we hypothesized that GS are metabolically dependent on intracellular cholesterol for survival. Statins inhibit HMG-CoA reductase (HMGCR), the first step in the sterol biosynthesis pathway, whose outputs include cholesterol, protein post-translational modifications (e.g., farnesyl, geranyl), steroid hormones, and coenzyme Q10 (CoQ10)^39, 40^ (**Figure 4I**). To determine the arm of the sterol biosynthesis pathway involved in mediating GS sensitivity to statins, we treated DIPG-007 GS *in vitro* with Pitavastatin alone or in combination with key intermediates of the sterol biosynthesis pathway, including mevalonate, farnesyl pyrophosphate (FPP), or geranylgeranyl pyrophosphate (GGPP) (**Figure 4I**). The results revealed that mevalonate, a rate limiting metabolite in the sterol biosynthesis pathway, protected cells from effects of Pitavastatin (**Figure 4J**). In addition, GGPP and FPP partially rescued Pitavastatin-induced loss of cell viability (**Figures 4K-L**).

We next investigated whether addition of exogenous cholesterol, an end product of the pathway, could similarly protect GS cells from Pitavastatin-induced cytotoxicity. Cholesterol conjugated to methyl-β-cyclodextrin to promote cell permeability was added to cells in combination with Pitavastatin. Here, we observed that cholesterol robustly rescued the loss of viability induced by Pitavastatin, indicating a dependency of GS on cholesterol for survival (**Figure 4M**). Similar observations were made using SF7761 and DIPG-XIII GS (**Supplementary Figures 8D-G; 8I-J**). CoQ_10_ acts as an electron shuttle between complexes II and III of the electron transport chain and is thus an important mediator of OXPHOS. However, CoQ_10_ media supplementation did not protect cells to the same extent as cholesterol or mevalonate. These results suggest that the cytotoxic effect of Pitavastatin is not the result of indirect inhibition of mitochondrial respiration via limiting CoQ_10_ biosynthesis (**Figure 4N; Supplementary Figures 8H,8K**).

### OPC-like GS exhibit decreased bioenergetic capacity and activity

To gain insights on why mitochondrial OXPHOS is a metabolic dependency in DIPG GS, we evaluated the bioenergetic capacity of untreated GS compared to DGC using the Seahorse extracellular flux analyzer. We monitored the oxygen consumption rate (OCR), which is an indicator of mitochondrial respiration. The results showed that the basal OCR was significantly lower in GS compared to DGCs in DIPG-007 and SF7761 but showed no significant difference in DIPG-XIII (**Figures 5A-B**). Further, challenge with the ATP synthase inhibitor, oligomycin, decreased respiration move severely in the GS. And, most strikingly, treatment with the mitochondrial membrane potential uncoupler FCCP, which facilitates maximal oxygen consumption in the mitochondria, revealed that the GS displayed markedly decreased OCR compared to DGCs (**Figure 5A**). Consistent with the lack of differential sensitivity of DIPG-XIII cells to OXPHOS inhibitors, there was no substantial difference in basal OCR and no response to Oligomycin or FCCP in GS vs. DGC (**Figures 5A-B**). Next, we assessed spare respiratory capacity (SRC), a measure of the difference between maximal oxygen consumption capacity and basal oxygen consumption in the mitochondria. SRC was similarly reduced in GS vs. DGC (**Figure 5C**). In alignment with our findings, cells with low SRC have been reported to be relatively proliferative and, low SRC is associated with stem-like cells, while SRC is elevated in the differentiated cells^41, 42^ (**Figure 1B**).

**Figure 5:**
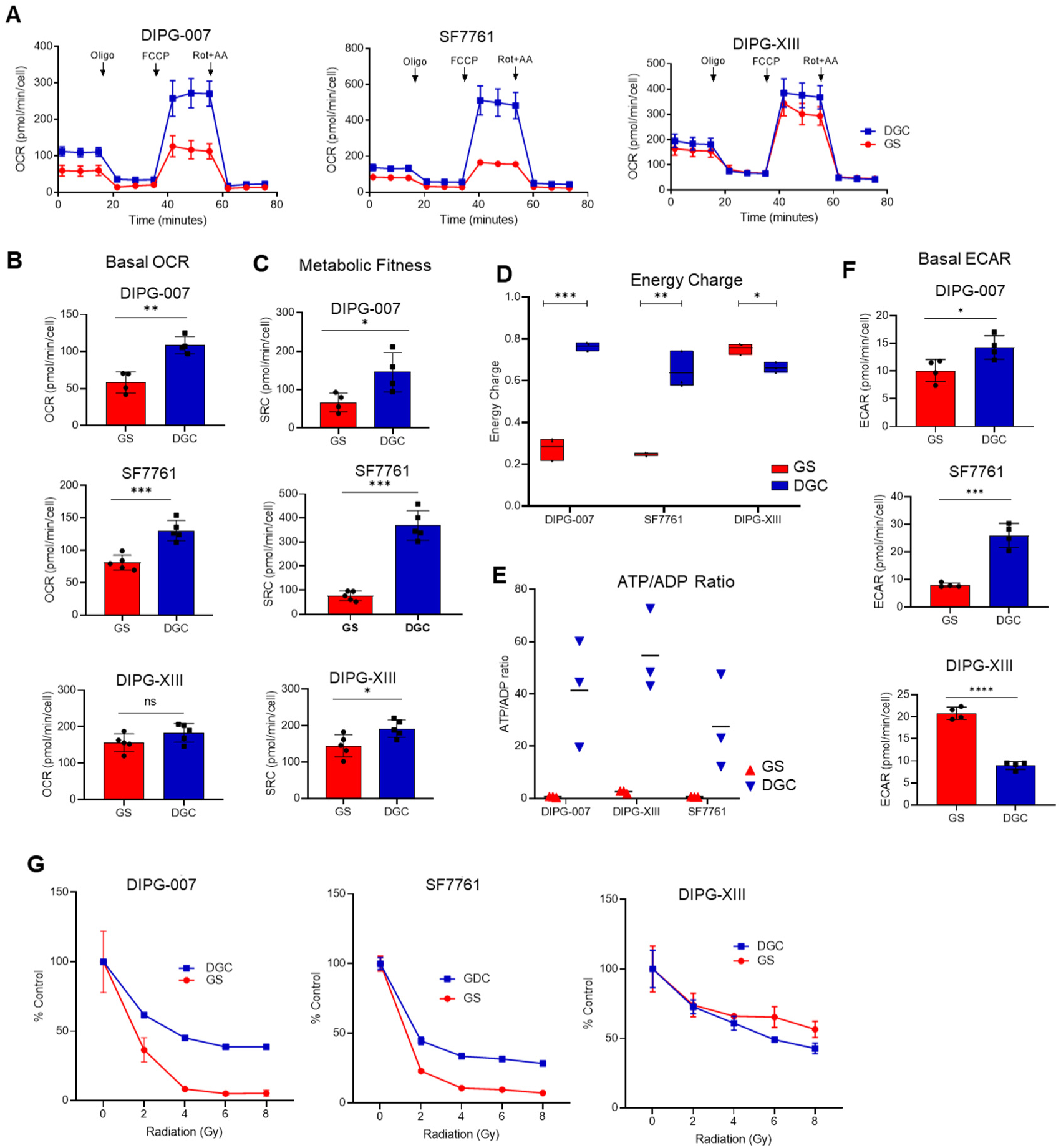
Bioenergetic properties of OPC-like GS and AS-like DGC. **A-C)** Bioenergetics analysis (Seahorse assay) of DIPG-007, SF7761, and DIPG-XIII GS and DGC. **A)** Oxygen consumption rate (OCR); **B)** Basal OCR, **C)** Spare respiratory capacity (SRC). Determination of **D)** energy charge calculated as [(ATP) + 0.5(ADP)]/ [(ATP) + (ADP) + (AMP)], and **E)** ATP/ADP ratios in GS and DGC across all three isogenic DIPG pairs. **F)** Extracellular acidification rate (ECAR) of DIPG-007, SF7761, and DIPG-XIII GS and DGC. Results expressed as mean ± SEM (ns = not significant; * p < 0.05; ** p < 0.01; *** p < 0.001; **** p < 0.0001). **G)** Dose-response curves of radiation-treated DIPG-007, SF7761, and DIPG-XIII GS and DGC. Cell viability assessed at 7 days post-treatment. Experiments were performed in technical triplicates and expressed percentage of control (untreated cells) and as mean ± SEM.

The seeming discrepancy between the upregulated mitochondrial OXPHOS gene signature (**Figure 1I**) and the decreased OCR and SRC parameters in the GS populations (**Figure 5B-C**) motivated us to take a more detailed look at the bioenergetic charge in our cultures. To this end, we interrogated our in-house metabolomics profiling dataset and determined both the adenylate energy charge and the ATP/ADP ratio of the cells. The adenylate energy charge (AEC) of a cell is an index of the energetic status of the cell that considers the differential intracellular levels of the adenylate pool, namely adenosine triphosphate (ATP), adenosine diphosphate (ADP), and adenosine monophosphate (AMP)^43, 44^. The AEC is calculated by applying the formula [(ATP) + 0.5(ADP)]/ [(ATP) + (ADP) + (AMP)], which yields values between 0 and 1 wherein normal cells remain in the 0.7 to 0.95 range^43, 44^. Assessment of the AEC in GS vs. DGC lines revealed that the tumorigenic DIPG-007 and SF7761 GS displayed markedly lower AEC values compared to their DGC counterparts (**Figure 5D**). This result is indicative of a consequent greater dependency of GS on ATP-generating pathway(s), chief among which is mitochondrial OXPHOS. Here again, modest but significant differences in the AEC values were observed for DIPG-XIII. This result is consistent with our observation that DIPG-XIII GS do not demonstrate the differential and selective sensitivity to OXPHOS inhibition.

Despite the variability in AEC among the isogenic DIPG pairs, the direct ratio of ATP to ADP revealed substantially lower levels in GS across all lines (**Figure 5E**). Thus, the low energy charge of GS indicates a DIPG metabolic state in which catabolic processes to regenerate ATP are limiting, which may provide an explanation for the therapeutic susceptibility to OXPHOS inhibition. Thus, we next analyzed glycolytic flux by measuring extracellular acidification rate (ECAR) using the Seahorse bioanalyzer. While upstream glycolytic pools were greatly enriched in DIPG-007 and SF7761 GS by metabolomics analysis (**Figure 2D**), ECAR was significantly more pronounced in their respective DGC counterparts (**Figure 5F**). These results suggest that DIPG-007 and SF7761 DGC can compensate for the inhibition of respiration through utilization of glycolysis, which the GS appear unable to do, potentially because of glycolytic stalling as reflected in the large metabolite pool sizes. In summary, these results reveal that OXPHOS inhibitor-sensitive GS have lower OCR, SRC, AEC, and ECAR. This suggests that DIPG-007 and SF7761 GS exist in a lower and more vulnerable bioenergetic state than their DGC counterparts, providing important insight into why OPC-like GS are highly sensitive to mitochondrial targeting.

Mitochondrial SRC correlates with the capacity of cells to respond or adapt to stress conditions (e.g. oxidative stress)^41^. We therefore hypothesized that the lower SRC in the GS would be reflected in an increased susceptibility to ionizing radiation, the mainstay therapy for DIPG and a well-established inducer of cytotoxic oxidative stress. To this end, we treated the DIPG isogenic cells with varying doses of radiation and evaluated cell viability after 7 days. With the exception of DIPG-XIII, the GS were markedly more sensitive to radiation treatment than DGC (**Figure 5G**).

### Cholesterol biosynthesis and OXPHOS inhibition decrease tumor burden and increase overall survival of DIPG tumor bearing mice

Statins are used to lower cholesterol and protect from cardiovascular disease and represent one of the most widely used drugs in the clinic, illustrating their safety and tolerability^45, 46^. Similarly, biguanides, which act through OXPHOS inhibition^47, 48^, are clinically deployed to reduce blood glucose in diabetes and have seen recent application in cancer trials, again illustrating the potential for rapid deployment in clinical trials for DIPG. Furthermore, studies have shown that biguanides can modestly transverse the blood brain barrier (BBB)^49, 50^, and some classes of statins display brain penetrance, depending on the pharmacophore, including Pitavastatin^51, 52^.

Thus, to evaluate the effects of OXPHOS inhibitors and statins on tumor growth and overall survival, we employed a preclinical orthotopic mouse model of DIPG in which bioluminescent DIPG-007 cells, grown under GS conditions, were stereotactically injected into the pons of immunodeficient mice. Tumor engraftment was confirmed via bioluminescent imaging (BLI) 3-weeks post tumor implantation and mice were randomized into four arms receiving vehicle, Pitavastatin (10mg/kg), Phenformin (50mg/kg) or a combination of both drugs, administered intraperitoneally (**Figure 6A**). These treatment doses were determined from an in-house dose-escalating tolerability study in which no signs of toxicity or weight loss were observed following administration of the drugs over a 2-week course. Notably, treatment with Pitavastatin or Phenformin resulted in the reduction in tumor volume based on BLI. Neither treatment adversely impacted mouse body weight (**Figures 6B-C)**. Consistent with our *in vitro* findings (**Supplementary Figures 7A-B)**, the combination of both drugs did not show improvement over single agent alone. In addition, treatment with Pitavastatin or Phenformin significantly extended the survival of DIPG-007 tumor-bearing mice, and here again, the combination did not provide additional benefit (**Figure 6D**). Collectively, single agent metabolic inhibitors showed promising results in providing survival benefits in this preclinical model of DIPG, demonstrating the potential utility of targeting cholesterol biogenesis and mitochondrial respiration in DIPG patients.

**Figure 6:**
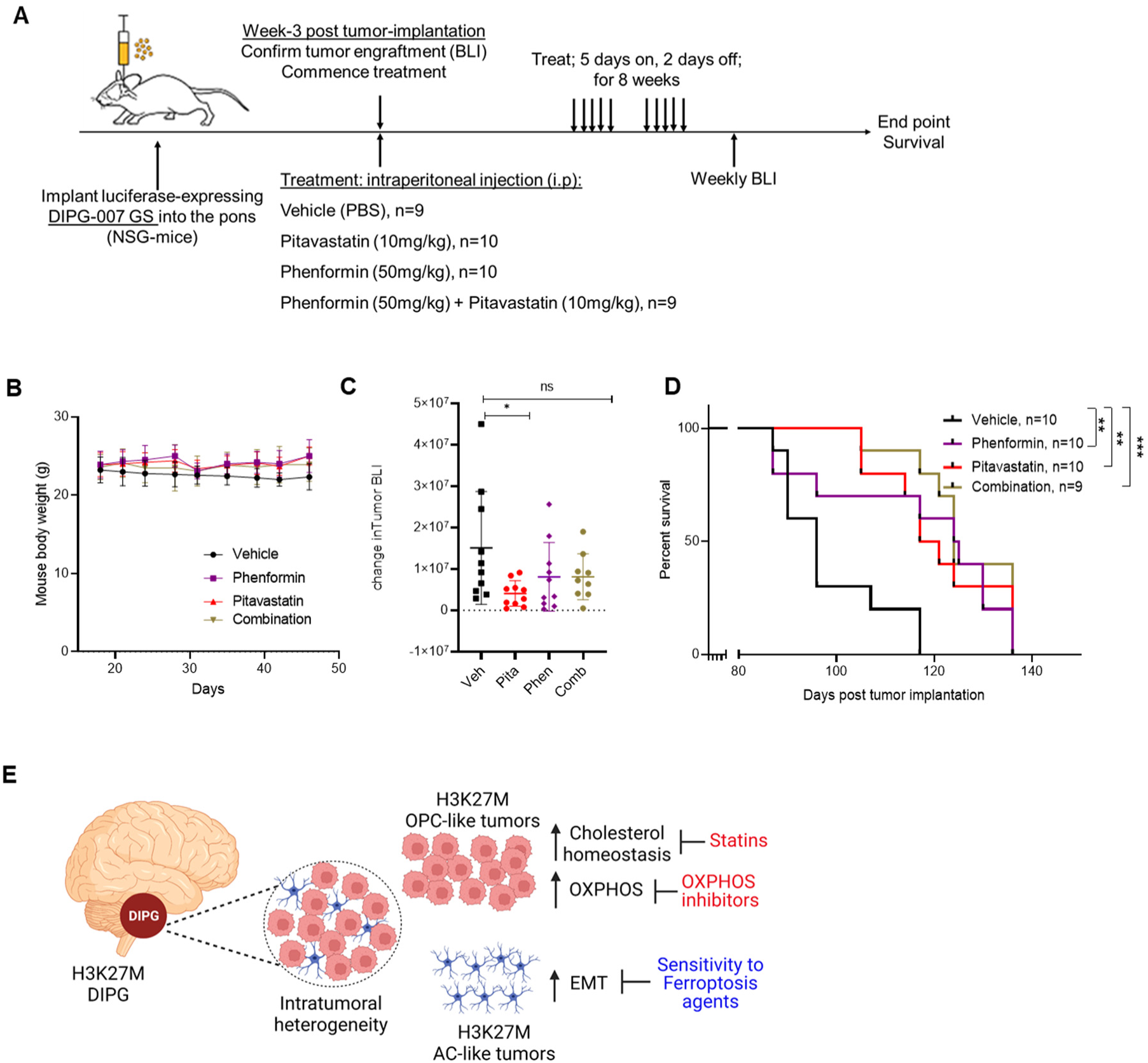
Statin and Complex I inhibitors prolong survival in an orthotopic model of DIPG. **A)** Schematic of *in vivo* experiment using DIPG-007 orthotopic xenograft model and intraperitoneal administration of Pitavastatin (10mg/kg; n=10), Phenformin (50mg/kg; n=10), or the combination (n=10) of both drugs at indicated doses or vehicle control (PBS; n=10). **B)** Body weight of DIPG-007 tumor-bearing NSG mice following treatment. **C)** Quantitation of change in tumor volume from bioluminescence imaging (BLI) over the course of week-3 to week-7 post-tumor implantation. **D)** End-point survival analyses of treatment and control tumor-bearing mice. **E)** Model of H3K27M DIPG intratumoral heterogeneity indicating specific vulnerabilities within OPC-like and AC-like tumor populations and their respective targeting strategies.

## DISCUSSION

H3K27M DIPGs are characterized by intratumoral heterogeneity comprising distinct tumor cell types, wherein the stem-like and tumor-initiating characteristics are driven by a population of less-differentiated OPC-like glioma cells while the more differentiated AC-like glioma cells represent a minority^12^. We demonstrated that this tumor heterogeneity can be modeled *in vitro* and is substantially recapitulated in isogenic DIPG GS and DGC, which are enriched for OPC-like and AC-like gene signatures, respectively.

By applying a systems biology-driven approach that encompassed transcriptomics, metabolomics, and bioenergetic analysis, we showed that the OPC-like and AC-like tumor phenotypes harbor distinct metabolic vulnerabilities. Compared to DGC, the GS populations showed higher levels of purine nucleotides. This finding is consistent with features of stem-like brain tumor-initiating cells described in adult glioblastoma (GBM), which upregulate purine synthetic intermediates to promote anabolic processes^32^. We also observed that GS exhibit increased intracellular levels of upstream glycolytic intermediates by metabolomics, though the rate of glycolysis (ECAR) was higher in DGC. These results suggest that glycolysis in GS may be stalled at the level of Enolase, and, moreover, that DGC are better positioned to circumvent the inhibition of mitochondrial respiration through enhanced glycolysis. Genotype-dependent analysis of metabolism in DIPG previously revealed elevated glycolysis in H3K27M gliomas compared to H3 wild-type tumors^18^. It will be important to test how the differentiation state interacts with the genotype to regulate glycolysis.

Along these lines, metabolites such as taurine, creatine, creatinine, uric acid, and hydroxyproline, which are reported to be associated with cellular differentiation of oligodendrocytes, cardiomyocytes, mesenchymal and adipocytes^33, 53–55^, were found to be upregulated in DGC. Indeed, taurine has been demonstrated to play a role in several biological processes, including the prevention of mitochondria damage, stabilization of OXPHOS in cardiomyocytes, and protection against endoplasmic reticulum (ER) stress. Creatine is involved in ATP buffering and enhancing mitochondria function^53^. Uric acid and hydroxy-proline have been linked to neuronal differentiation of mesenchymal stem cells^54^ and the differentiation of retinal pericytes to adipocytes^55^, respectively. These results suggest that these metabolites are pertinent to cellular differentiation processes, irrespective of the cell of origin.

Our transcriptomics analysis revealed AC-like DGC exhibited an enhanced mesenchymal phenotype. Based on this insight, we demonstrated that DGC were more sensitive to agents that promote ferroptosis. Conversely, OPC-like GS cells, whose gene signature correlated with higher disease aggressiveness and decreased overall survival in patients, upregulated cholesterol metabolism and mitochondrial OXHPHOS. Targeting these pathways with Phenformin and Metformin (mitochondria complex I inhibitors) or statins (sterol biosynthesis inhibitor) resulted in selective killing of GS compared to DGC *in vitro*. As a proof of principle, we also demonstrated considerable *in vivo* activity of these metabolic inhibitors in an orthotopic mouse model of DIPG (**Figure 6E**).

In DIPG-007, SF7761, and murine PPK isogenic pairs, the GS populations could be selectively targeted by inhibiting OXPHOS. In contrast, DIPG-XIII isogenic pair showed a limited differential phenotype to OXPHOS inhibition. Therefore, future studies with these models could help to determine predictive biomarkers of sensitivity to OXPHOS targeting. It is conceivable that the limited differential phenotype between DIPG-XIII GS and DGC in outcomes such as cell proliferation, purine nucleotide pools, TCA cycle metabolites, OCR, SRC, energy charge, sensitivity to radiation, and sensitivity to OXPHOS inhibitors may result from oncogenic signaling related to *MYC* and *EGFR*. Indeed, the greater than two-fold upregulation of *MYC* and *EGFR* seen in DIPG007 and SF7761 GS, in comparison to their GDC counterparts, was not similarly observed in DIPG-XIII GS vs. DGC. Along these lines, a question that merits future investigation is whether specific oncogenic signaling pathway(s) or transcription factor(s) operating in distinct tumor subpopulations direct metabolic reprogramming. For instance, MYC and EGFR have been reported to be critical for maintenance of the brain tumor-initiating cells in adult glioblastoma^56, 57^.

The concentration of cholesterol is highest in the brain, at approximately 20% of total body cholesterol^58^. In addition, the majority of brain cholesterol results from *de novo* synthesis, rather than uptake from circulation or peripheral tissues^59^. These results may provide mechanistic insight into the dependence of stem-like GS on cholesterol, and not on other outputs of the sterol biosynthesis pathway. Further, astrocytes are known to be the predominant producers and suppliers of cholesterol to other cells in the brain, including cancer cells^60^. Indeed, a dependency on cholesterol and the liver X receptors (LXR) axis as well as lanosterol synthase has been reported in brain tumors^61, 62^. Our study, therefore, adds to the growing evidence of a metabolic dependency of brain tumors on cholesterol and specifically presents cholesterol targeting as a novel therapeutic inroad for stem-like and tumorigenic H3K27M midline gliomas.

Targeting DIPG via OXPHOS and cholesterol inhibition is a promising strategy in that the inhibitors of these pathways are clinically approved drugs and have been evaluated as chemo-sensitization agents in cancer clinical trials^46, 47^. Moreover, a number of statins are known to penetrate the blood brain barrier^51, 52^. In addition, Metformin, a biguanide and analog of phenformin, has been used in the clinic for several decades, and it is currently being tested in several cancer clinical trials as a chemo-adjuvant. Importantly, it too displays some degree of brain penetrantance^50^. Therefore, targeting mitochondria OXPHOS and cholesterol biosynthesis could potentially have immediate clinical utility for DIPG patients. Lastly, given the limited combinatorial activity of OXPHOS inhibitors and statins, future studies will be required to test efficacy alongside ionizing radiation therapy.

Metabolic dependencies have been investigated in H3K27M gliomas in comparison to H3WT tumors or normal brain tissue, and these studies have revealed dependencies on glucose and glutamine metabolism^18^. To the best of our knowledge, this is the first study to investigate differentiation-state dependent metabolic vulnerabilities in H3K27M DIPGs. Our study, therefore, adds to the growing body of work on DIPG metabolism and presents novel actionable metabolic vulnerabilities that can be leveraged to develop new treatment options for this devastating disease. Indeed, the findings from this study are significant in that they provide a framework for future investigations that could, by extension, have broad implication in the rational design of precision treatment approaches for H3K27M DIPG patients based on tumor composition and abundance of specific tumor cell-types.

## METHODS

### Cell lines and culture conditions

All cell lines used were routinely tested for mycoplasma and were validated by STR profiling. HSJD-DIPG-007 (referred to as DIPG-007, H3.3K27M) was obtained from Dr. Rintaro Hashizume, Northwestern University; RRID: CVCL_VU70. SU-DIPG-XIII (referred to as DIPG-XIII, H3.3K27M) was obtained from Dr. Michelle Monje, Stanford University; RRID: CVCL_IT41. SF188 (H3WT) and normal human astrocytes (NHA, H3WT) were obtained from Dr. Craig B. Thompson, Memorial Sloan Kettering Cancer Center; RRID: CVCL_6948. SF7761 (H3.3K27M) was purchased from Millipore Sigma (Cat.no. SCC126). All cells were cultured in a humidified incubator at 37°C and 5% CO_2_. DIPG007 and DIPG-XIII gliomaspheres were cultured in base media containing equal parts Neurobasal A and DMEM/F12 with added HEPES buffer (10 mM), MEM sodium pyruvate solution (1mM), MEM Non-Essential Amino Acids (1X), GlutaMAX-I Supplement (1X), Antibiotic-Antimycotic (1X), and supplemented with fresh B27 (without vitamin A), heparin (2 μg/mL), human-EGF (20 ng/mL), human-bFGF (20 ng/mL), PDGF-AA (10 ng/mL), and PDGF-BB (10 ng/mL). SF7761 gliomaspheres were cultured in a base media containing Neurobasal A with added N2, B27, L-glutamine (2 mM), Pen/strep (1X), and supplemented with fresh heparin (2 μg/mL), human-EGF (20 ng/mL), human-bFGF (20 ng/mL), and BSA (45 ng/ml).

PPK cell line was received as a gift from Dr. Carl Koschmann, University of Michigan. PPK cells were generated as an In Utero Electroporation (IUE) murine model of H3K27M glioma. IUE was performed using sterile technique on isoflurane/oxygen-anesthetized pregnant C57BL/6 or CD1 females at E13.5. Tumors were generated with lateral ventricle (forebrain) introduction of plasmids: (1) PB-CAG-DNp53-Ires-Luciferase (dominant negative T<u>P</u>53), (2) PB-CAG-PdgfraD824V-Ires-eGFP (<u>P</u>DGFRA D842V), and (3) PB-CAG-H3.3 <u>K</u>27M-Ires-eGFP (H3K27M), and therefore referred to as “PPK” model^38^. PPK gliomaspheres were cultured in base media containing Neurobasal-A medium (1X), MEM sodium pyruvate solution (1mM), MEM Non-Essential Amino Acids (1X), L-glutamine (2 mM), Antibiotic-Antimycotic (1X), and supplemented with fresh B27, N2, heparin (2 μg/mL), EGF (20 ng/mL), FGF (20 ng/mL).

Differentiated human and murine H3K27M glioma cells were generated by dissociating the respective gliomaspheres into single cells with Accutase (Stemcell Technologies; # 07920) and subsequently cultured and maintained in the respective base media supplemented with 10% FBS for 14 days to generate a monolayer adherent culture. SF188 and NHA cells were cultured in DMEM supplemented with FBS (10%), L-glutamine (2 mM), and Pen/strep (1X).

### Growth Assay

Proliferation was calculated by automated counting of cells. Differentiated glioma cells or gliomaspheres were dissociated into single cells with trypsin or Accutase, respectively. Following this, 200,000 cells were plated in 60mm dishes, and cultured in their respective growth media for up to 12 days with media replenished every 3 days. At the indicated time, cells were dissociated using trypsin (differentiated cells) or Accutase (gliomaspheres) and enumerated using the Countess II FL Automated Cell Counter.

### Drug treatment and viability assay

The following compounds used in this study were purchased from Cayman Chemicals: (1S,3R)- RSL3 (RSL3, # 19288), Ferrostatin-1 (#17729), z-vad-FMK (#14463), Necrosulfonamide (#20844), Bafilomycin A-1 (#11038), Metformin (#13118), Phenformin (#14997), IACS-010759 (#25867), Atorvastatin (#10493), Fluvastatin (#10010334), Pitavastatin (#15414), Mevalonate (#20348), Farnesyl Pyrophosphate (#63250), Geranylgeranyl Pyrophosphate (#63330), and Coenzyme Q10 (#11506). Cholesterol-Water Soluble (Cholesterol–methyl-β-cyclodextrin, #C4951) and Trolox (#238813) were purchased from Millipore Sigma. Equal numbers of isogenic DIPG-007, SF7761, DIPG-XIII GS and DGCs were plated in white opaque 96-well plate at 2,000-3,000 cells per well and incubated overnight. Cells were treated with compounds at the indicated concentrations and length of time as described in the figure legend. At end point, an equal volume of Cell Titer Glo (2.0) reagent (Promega, G9243) or 3D-Cell Titer Glo (Promega, G9683) was added to each well and viability assessed according to manufacturer’s protocol. Luminescence was detected and measured using a SpectraMax M3 plate reader and data analyzed with GraphPad Prism 8 software.

### Detection of Lipid ROS

To assess levels of lipid ROS in cells, 200,000 DGCs were plated in a 6-well plate overnight and treated with the indicated compounds. At end point, cells were washed twice with PBS and stained for 20 minutes with 2 µM C11-BODIPY (Invitrogen, D3861) in a phenol red-free media. Following staining, cells were washed twice with PBS and dissociated to singles cells with trypsin. The cells were then transferred to round-bottom 96-well plates on ice, co-stained with Sytox-blue (Invitrogen, S34857) to distinguish viable cells, and analyzed on a ZE5 Cell analyzer (Bio-Rad). C11-BODIPY signal was captured with the FITC channel. Analysis of data was performed using FlowJo v.10 software.

### Seahorse Bioenergetics Assay

The cellular bioenergetic state was analyzed using a Seahorse XF-96 Extracellular Flux Analyzer (Agilent). The sensor cartridges were incubated in dH_2_O overnight, and on the day of the assay, the cartridges were hydrated in XF calibrant (Agilent) for 1 hour in a non-CO_2_ incubator at 37^0^C. The hydrated cartridges were loaded with oligomycin (1μM), FCCP (1μM), rotenone (0.1μM), and antimycin A (1μM) to perform the Mito stress test. Concurrently, 96-well Seahorse cell culture plates were coated overnight with laminin, and on the day of the assay, dissociated DIPG GS were washed and resuspended in a Seahorse XF RPMI media (Agilent;103576) supplemented with XF Glutamine (Agilent;103579), 10mM glucose, MEM sodium pyruvate solution (1mM), MEM Non-Essential Amino Acids (1X), human-EGF (20 ng/mL), human-bFGF (20 ng/mL), PDGF-AA (10 ng/mL), and PDGF-BB (10 ng/mL), while dissociated DGCs were washed and resuspended in a Seahorse XF RPMI media (Agilent;103576) supplemented with XF Glutamine (Agilent;103579), 10mM glucose, MEM sodium pyruvate solution (1mM), and MEM Non-Essential Amino Acids (1X). DGC and GS single cells (150,000 to 200,000 cells)were seeded on laminin-coated plates and allowed to equilibrate for 30 minutes in a non-CO_2_ incubator at 37°C. Data were then acquired on the Seahorse analyzer. Following data acquisition, measurements were normalized based on cell number using the CyQuant Cell Proliferation Assay (Invitrogen). For the Mito stress test, the basal oxygen consumption rate (basal OCR) was determined based on basal OCR measurements taken prior to addition of inhibitors. The spare respiratory capacity (SRC) was determined by subtracting basal OCR from maximal OCR measurements. Seahorse analysis was performed using the Wave 2.3 software.

### Metabolomics

#### Metabolite extraction

To generate intracellular metabolite fractions, an equal number of GS and DGCs were cultured in 6-well plates for 36 hours. Next the growth media was removed, cells were lysed with ice-cold 80% methanol on dry ice for 20 minutes. Lysates were collected and clarified by centrifugation. The metabolite load of intracellular fractions was normalized to protein content of parallel samples and these volumes were then lyophilized in a SpeedVac. Dried metabolite pellets were resuspended in 50:50 mixture of HPLC-grade methanol:dH20 and subjected to metabolomics analysis.

#### LC/MS-based Metabolomics

LC/MS-based Metabolomics was performed on an Agilent 1290 Infinity II LC-coupled to a 6470 Triple Quadrupole (QqQ) tandem mass spectrometer (MS/MS), as previously described^63^. Briefly, Agilent Masshunter Workstation Software LC/MS Data Acquisition for 6400 Series Triple Quadrupole MS with Version B.08.02 was used for compound optimization, calibration, and data acquisition. The QqQ data were pre-processed with Agilent MassHunter Workstation QqQ Quantitative Analysis Software (B0700). For post-sample normalization, total metabolite ion currents in each sample were summed, and this value was applied proportionally to the sample values for proper comparisons, statistical analyses, and visualizations among metabolites. Two-tailed t-test with a significance threshold level of 0.05 was applied to determine statistical significance between conditions. Graphs were generated using GraphPad Prism 8.0 software. Heatmaps were generated and data clustered using Morpheus Matrix Visualization and analysis tool (https://software.broadinstitute.org/morpheus). Pathway analyses were conducted using MetaboAnalyst (https://www.metaboanalyst.ca).

### Energy Charge Calculation

For each sample, the ion current from the LC/MS analysis for adenosine triphosphate (ATP), adenosine diphosphate (ADP), and adenosine monophosphate (AMP) levels were enumerated, and the adenylate energy charge (AEC) was calculated by applying the formula [(ATP) + 0.5(ADP)]/ [(ATP) + (ADP) + (AMP)]. The ratio of ATP/ADP was evaluated by directly determining the ratio of ATP to ADP metabolite levels in each cell line.

### RNA sequencing

Total RNA was extracted from DIPG-007, SF7761, and DIPG-XIII GS and DGCs using the RNeasy Mini Kit (Qiagen) according to the manufacturer’s instructions. Strand-specific, poly- A+ libraries were prepared using NEBNext Ultra II Directional RNA Library Prep Kit (New England Biolab, E7760L), the Poly(A) mRNA Magnetic Isolation Module (New England Biolab, E7490L), and NEBNext Multiplex Oligos for Illumina Unique Dual (New England Biolab, E6440L). Sequencing was performed on the NovaSeq-6000 (Illumina), yielding 150-base, paired-end reads. Library preparation and sequencing were performed by the University of Michigan Advanced Genomics Core (Ann Arbor, MI).

The reads were trimmed using Trimmomatic v0.36^64^ and the library qualities were assessed using FastqQC v0.11 for trimmed reads (https://www.bioinformatics.babraham.ac.uk/projects/fastqc/). RSEM v1.3.1 and STAR v2.5.2a were used to generate paired-end alignments and counts^65, 66^.

Differential gene expression analysis was performed using DESeq2 v1.26.0 and the *apeglm* shrinkage estimator was used to adjust log_2_ fold-changes^67^. Normalized counts were obtained DESeq2 (default method; median of ratios). Differentially expressed genes were defined as having adjusted p-value < 0.05 and fold change > 1.5 (up or down). Variance stabilized transform (VST) gene counts were used in principal component analysis to identify the major sources of variance and evaluate the similarity of replicates. The reference sequence hg38 (GRCh38) and annotations, including gene IDs, were obtained from GENCODE v29. Differentially expressed genes were analyzed using GSEA using the HALLMARK gene sets.

### DIPG and DMGs dataset analysis

The human DIPG and DMG dataset was mined from Mackay et al ^28^. Expression levels of NS related genes in 76 H3K27M diffuse midline gliomas were segregated into high vs. low gene expression categories using unbiased K-means clustering (K=2 to assign two groups). Kaplan-Meier analysis was then performed between high (defined as upper quartile) vs. low (all remaining samples) tumors to determine differences in overall survival. Data were analyzed by the Log rank test.

### Western Blot

To assess protein levels of OLIG2, DIPG-007, SF7761, DIPG-XIII GS and GDC were lysed in RIPA buffer (Sigma, #R0278) containing protease (Roche, #04693132001) and phosphatase (Sigma, #P5726) inhibitors. To determine changes in apoptosis markers, DIPG-007 DGC and GS were treated as described in figure legends and lysed. Protein concentrations from whole cell lysates were determined using Pierce BCA Protein Assay kit (#23227), according to the manufacturer’s protocol. Equal amounts of protein were subjected to separation on SDS-PAGE and transferred to a methanol-activated PVDF membrane. Membranes were blocked with 5% milk in TBST (Tris-buffered saline containing 0.1% Tween 20) followed by incubation with primary antibodies diluted in 5% milk or BSA TBST at 4°C overnight. The following primary antibodies and dilutions were used: H3 histone (Cell Signaling Technology, CST #4499S; 1:1000), H3K27M (CST #74829S; 1:1000), Tri-methyl-Histone H3 (K27) (CST #9733S; 1:1000), Olig2 (CST #65915; 1:1000), PARP (CST #9542S; 1:1000), HSP90 (CST #4874S; 1:10,000) and Vinculin (CST #13901S; 1;10,000). Following primary antibody incubation, the membranes were washed 3 times with TBST and incubated with species-appropriate secondary antibodies conjugated to horse radish peroxidase (HRP) at 1:10,000 dilution for 1 hour at room temperature. Membranes were then washed 5X with TBST and chemiluminescence was detected using Clarity (Biorad, #1705060) or Clarity Max (#1705062) ECL substrate. The signal was captured with a Biorad Chemidoc imager and analyzed using Image Lab software.

### Mouse studies

Animal experiments were performed after approval from the University of Michigan Committee on Use and Care of Animals and were conducted as per NIH guidelines for animal welfare. All animal procedures were approved by Institutional Animal Care & Use Committee (IACUC) at the University of Michigan (IACUC approval # PRO00008865). Animals were housed and cared for according to standard guidelines with free access to food and water. All experiments were performed on NOD-SCID-IL2R gamma chain-deficient (NSG) mice that were 8–10 weeks old, with males and females used equally. Animals, including littermates of the same sex, were randomly assigned to control or treatment conditions.

### *In vivo* xenograft tumor studies

Luciferase-expressing DIPG-007 GS (400,000 cells) suspended in 2μl PBS were injected into the pons to establish orthotopic xenografts under anesthesia, as follows. NSG mice were anesthetized with 75mg/kg dexmedetomidine and 0.25mg/kg ketamine by intraperitoneal injection. Carprofen (5mg/kg) was used for analgesia. Mice were mounted on a stereotaxic device. A small sagittal incision was made using a scalpel and a small hole was created using a micro drill at 1.0 mm posterior and 0.8 mm lateral left from lambda. A sterile Hamilton syringe was used to inject cells. Half of the cells were injected at 5 mm depth from the inner base of the skull and the remaining cells were injected after 0.5 mm retraction in order to implant cells into the pontine tegmentum. After surgery, 1mg/kg atipamezole solution was intraperitoneally injected for anesthesia reversal. Tumor engraftment was confirmed by bioluminescence imaging. Treatment commenced at 3 weeks post tumor implantation. The mice were randomized into 4 groups receiving either vehicle (PBS), Pitavastatin (10mg/kg), Phenformin (50mg/kg), or combination Pitavastatin (10mg/kg) and Phenformin (50mg/kg). The drugs were administered intraperitoneally using a 5-day on/2-day off course for 9 weeks. Tumor size was measured using bioluminescent imaging (IVIS) up to 10-weeks post-implantation, at which mice were then monitored for end-point survival.

### Statistical Analysis

Statistical analyses were performed using GraphPad Prism 8 (Graph Pad Software Inc). Two-group comparisons were analyzed using the unpaired two-tailed Student’s t test. Error bars represent mean ± standard deviation, unless noted otherwise, and the significance annotations are indicated in each figure. A p-value < 0.05 was considered statistically significant. The number and type of experimental replicates as well as the explanation of significant values are indicated in the figure legends. All experiments were repeated at least twice.

### Data availability

RNAseq data are deposited at the GEO public repository using the following identifier, GSE197145. Raw metabolomics data are provided as Supplemental Table 1. All other datasets generated and or analyzed during the study are available from the corresponding author on reasonable request.

## Acknowledgements

This work was funded by a joint Defeat DIPG and ChadTough Foundation fellowship award (NEM); Alex’s Lemonade Stand Foundation (SV and CAL); and the University of Michigan Pediatric Brain Tumor Initiative (CK, SV, and CAL). Metabolomics studies performed at the University of Michigan were supported by the Charles Woodson Research Fund and the UM Pediatric Brain Tumor Initiative. Research reported in this publication was also supported by the National Cancer Institutes of Health under Award Number P30CA046592 using the following Cancer Center Shared Resource(s): Flow Cytometry Core, Tissue and Molecular Pathology Core. Schematics and models were created using Biorender.com.

## Author contributions

NEM and CAL conceived of and designed this study. NEM, CJK, SV and CAL planned and guided the research. NEM, ALM, CC, JKT, HSH, PS, ZCN, MS, SRS, DM, BC, LZ, BM, ZZ performed experiments, analyzed, and interpreted data. NEM, DRW, LF, SA, CJK, SV, and CAL were involved in the conceptual design of experiments and proofreading of manuscripts. NEM and CAL wrote the manuscript. SV and CAL supervised the work carried out in this study.

---

**Supplementary Figure 1:**
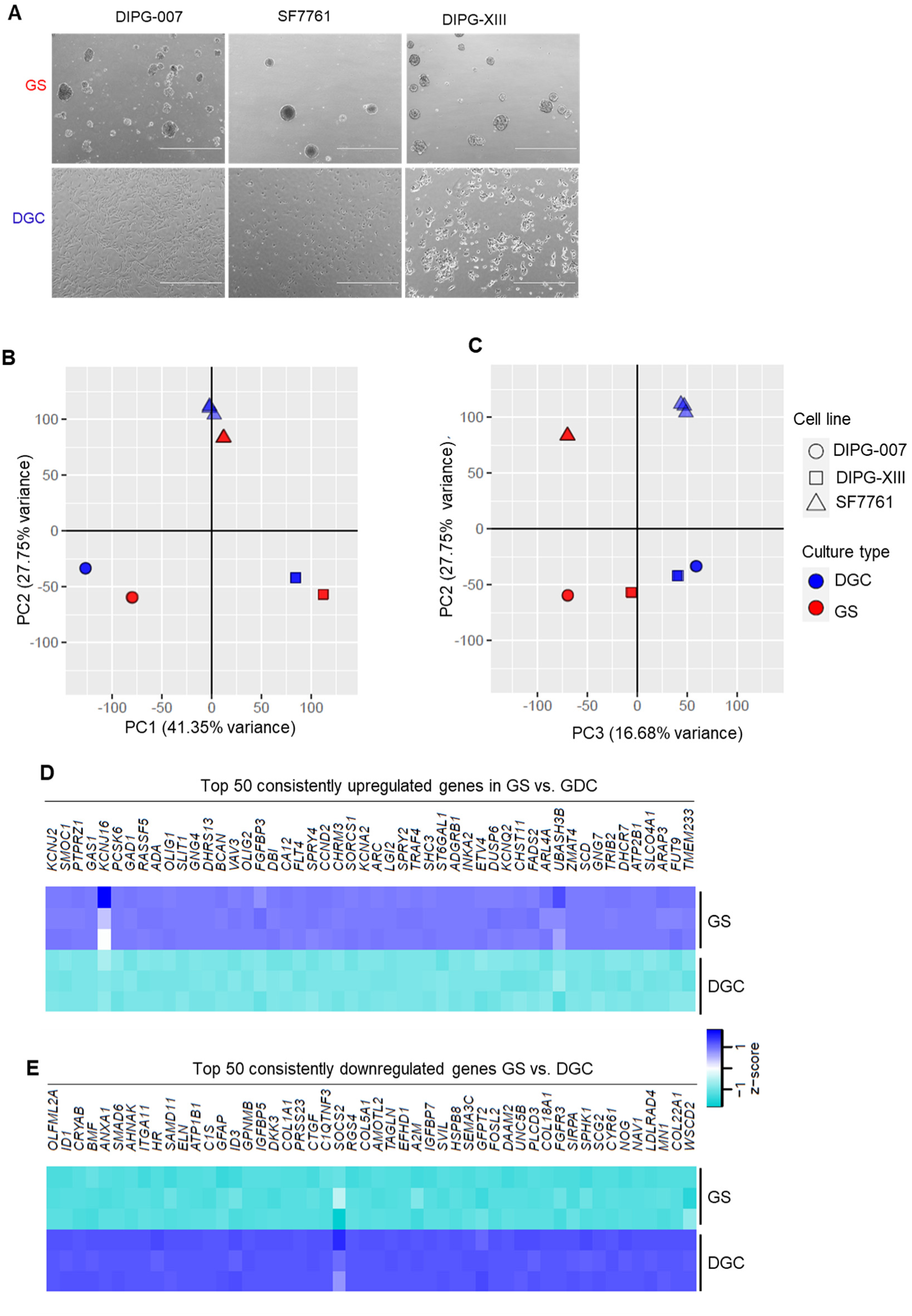
Characterization and transcriptomics analysis of isogenic DIPG models. **A)** Representative phase contrast microscopy images of patient-derived DIPG cell lines: DIPG-007, SF7761, and DIPG-XIII, showing morphological differences between isogenic gliomaspheres (GS) and differentiated glioma cells (DGC); scale bars indicate 1000 um. **B, C)** Principal component analysis of RNA-seq data. **D, E)** Heatmaps indicating the top 50 consistently and significantly upregulated **D)** and downregulated **E)** genes common to all three isogenic GS vs. DGC isogenic lines (adjusted p-value < 0.0001).

**Supplementary Figure 2:**
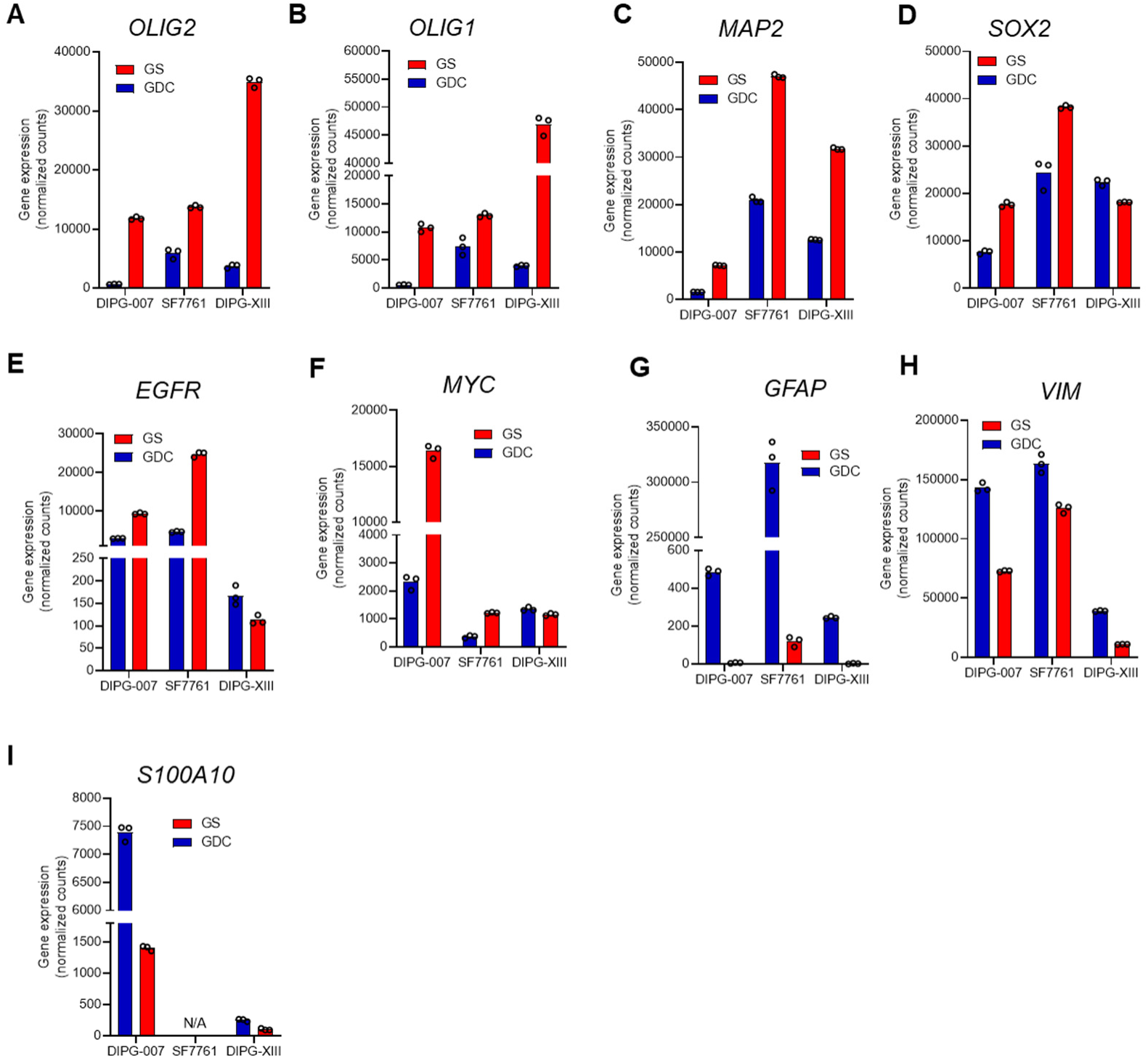
Transcriptomics analysis of gene markers in isogenic DIPG models. Relative gene expression of DIPG stemness and differentiation markers in GS vs. DGC across DIPG-007, SF7761, and DIPG-XIII. **A)** *OLIG2*, **B)** *OLIG1*, **C)** *MAP2*, **D)** *SOX2*, **E)** *EGFR*, **F)** *MYC*, **G)** *GFAP*, **H)** *VIM*, **K)** *S100A10*. Gene expression normalized counts plotted from the bulk transcriptomics analysis with technical replicates.

**Supplementary Fig. 3:**
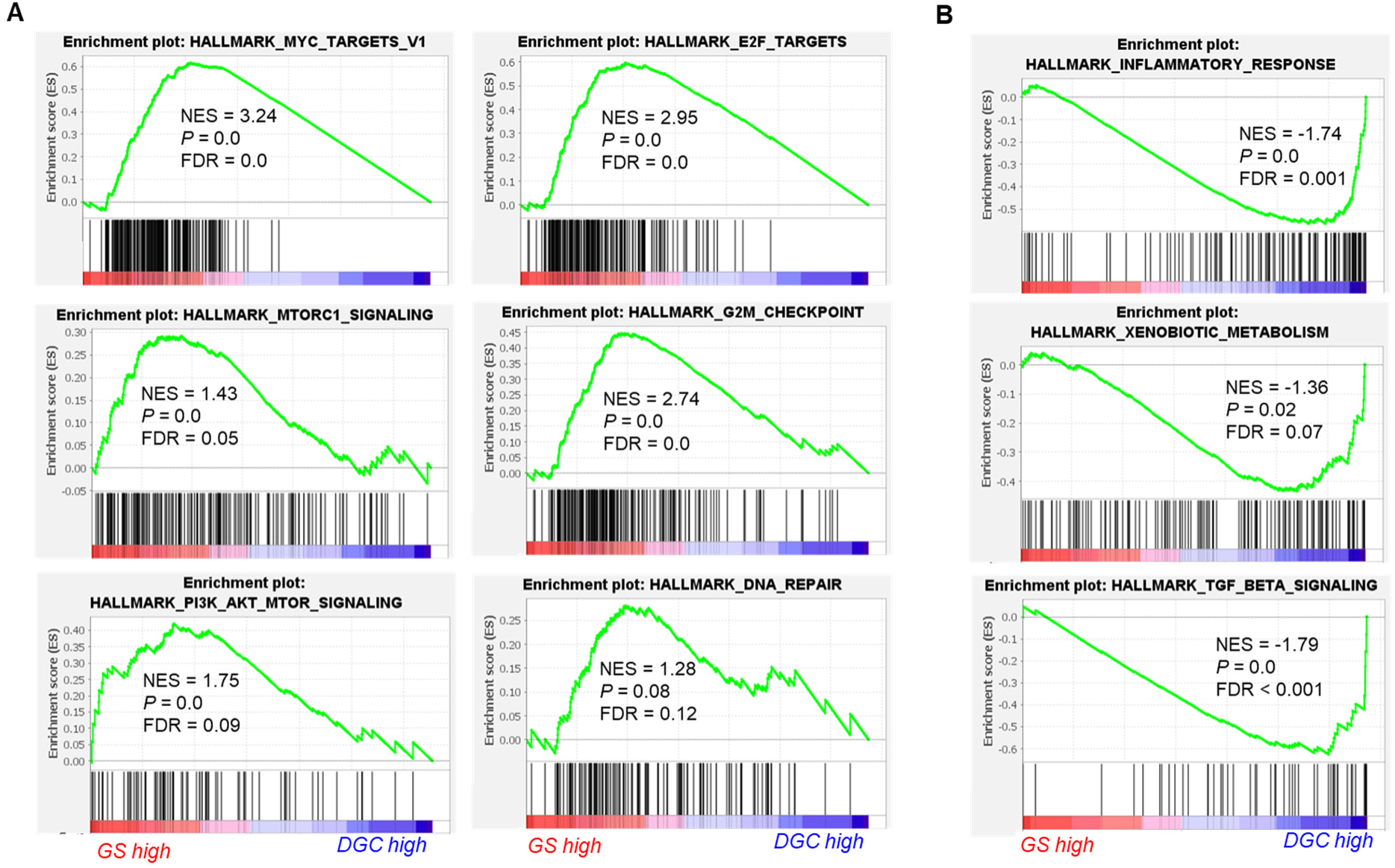
Gene set enrichment pathway analysis (GSEA) of isogenic DIPG cells. Representative gene set enrichment analysis (GSEA) indicating pathways that are **A)** upregulated, and **B)** downregulated in DIPG GS vs. DGC across all three isogenic lines. GSEA plots show enrichment scores and include values for normalized enrichment score (NES), nominal p value (*P*), and false discovery rate (FDR) q value.

**Supplementary Fig. 4:**
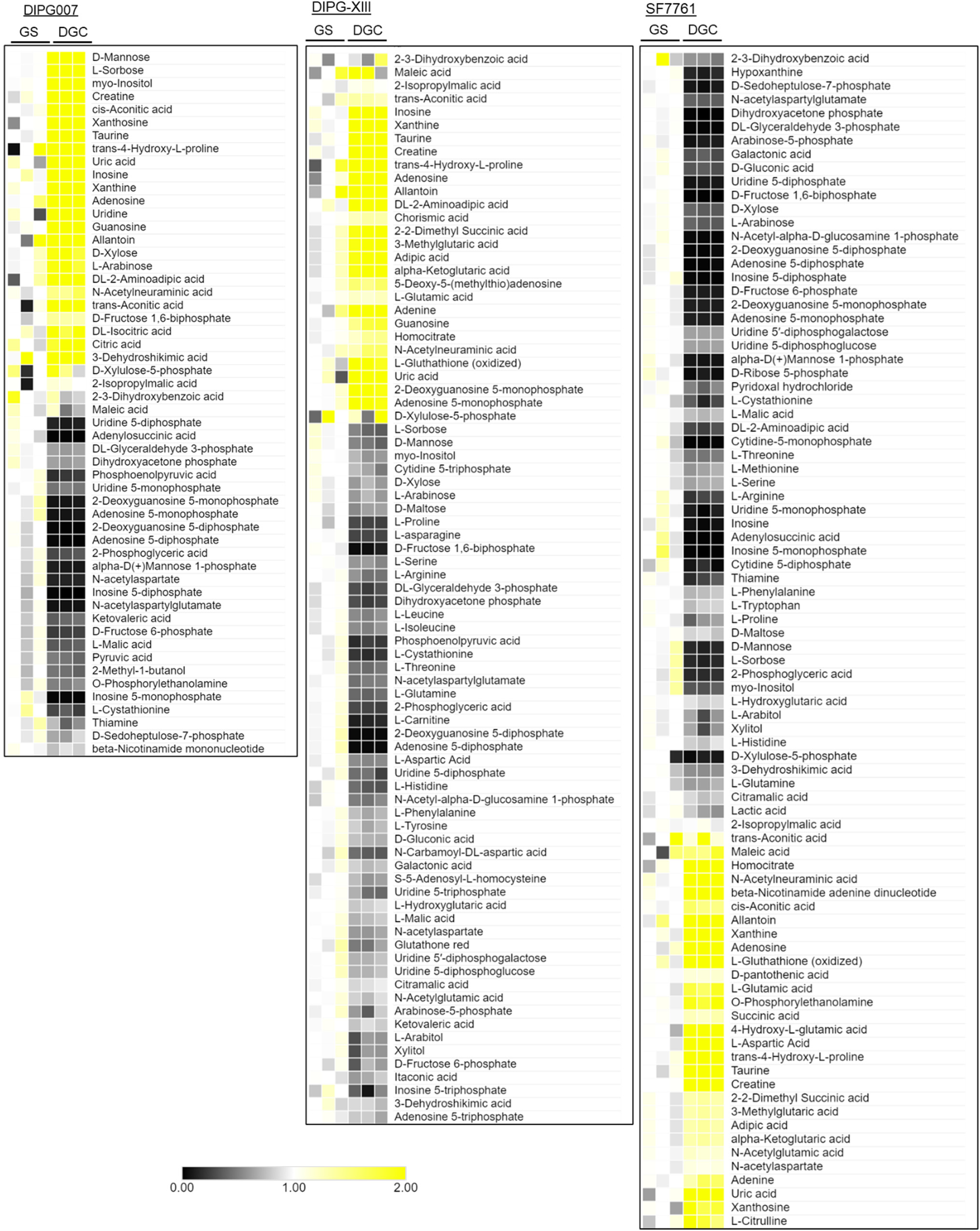
Steady-state metabolomics profiling of isogenic DIPG cells. **A)** Heatmaps showing significantly altered (p < 0.01) and differential abundance of metabolites seen in the three isogenic lines. Columns represent biological replicates; rows represent median normalized metabolites within each pair of isogenic lines.

**Supplementary Fig. 5:**
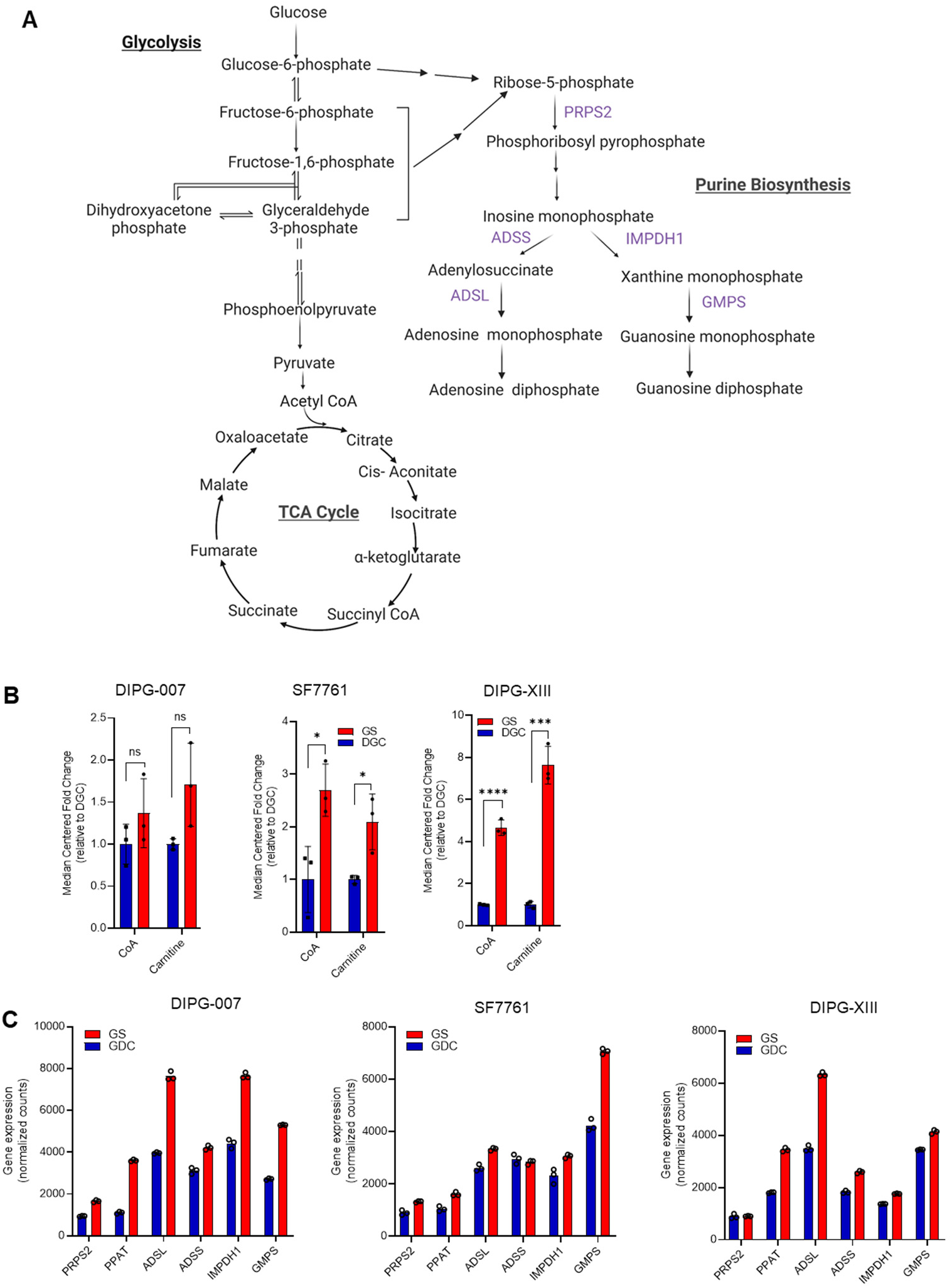
Metabolic profile of isogenic DIPG cells. **A)** Schematic of central metabolic pathways highlighting connections in glycolysis, TCA cycle, and purine biosynthetic pathways. **B)** Levels of CoA and Carnitine in DIPG-007, SF7761, and DIPG-XIII GS vs. DGC population. Metabolites levels expressed as median centered fold change of GS related to DGC levels across the three isogenic lines (ns = not significant; * p < 0.05; ** p < 0.01; *** p < 0.001; **** p < 0.0001). Error bars represent mean ± SD. **C)** Relative expression of genes (bulk RNA-seq data) encoding enzymes involved in purine biosynthesis in DIPG-007, SF7761, and DIPG-XIII isogenic GS and DGC counterparts.

**Supplementary Figure 6:**
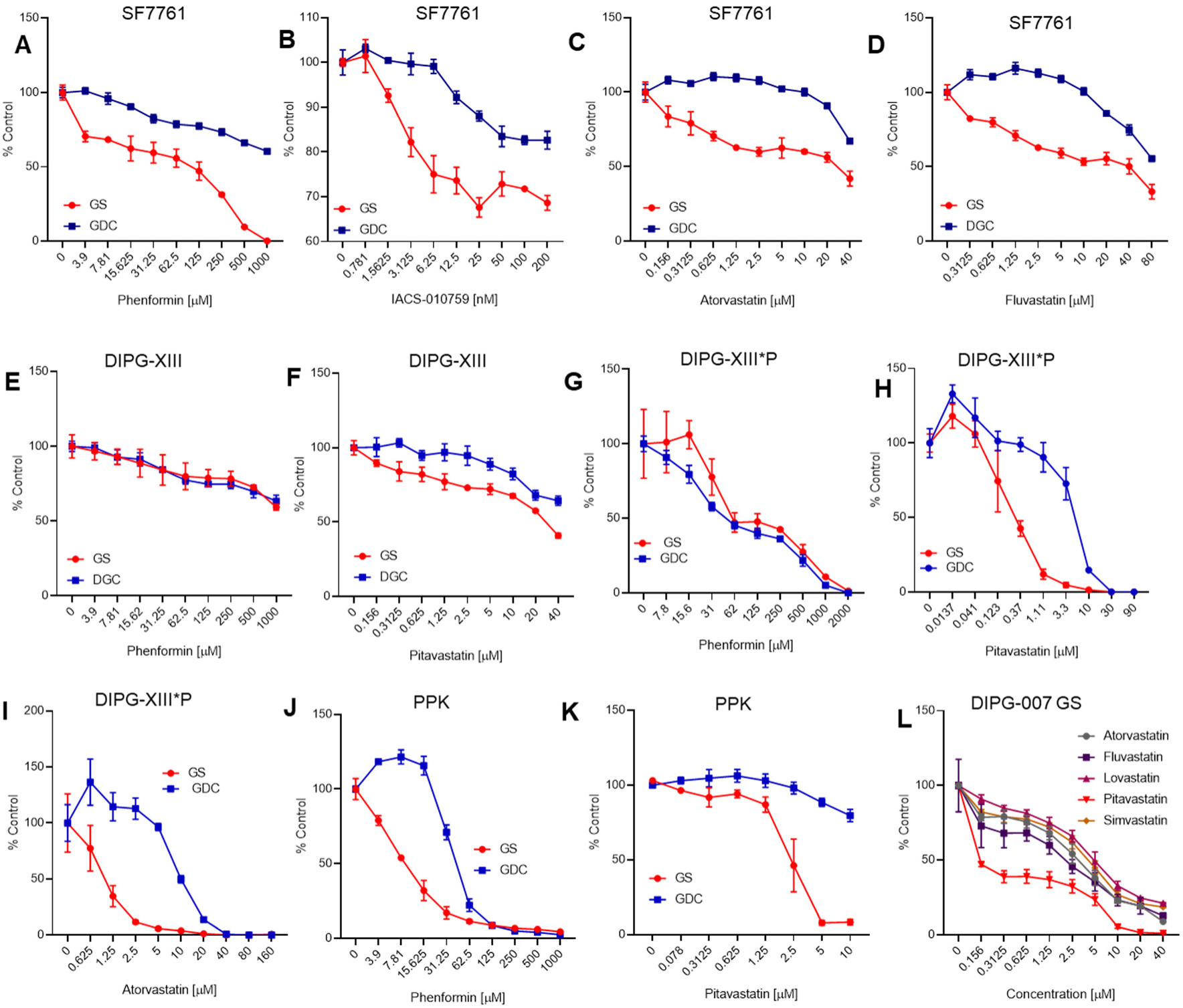
In vitro targeting of isogenic DIPG cultures with OXPHOS inhibitors and Statins. **A-D)** Dose response curves of SF7761 GS and DGC treated with **A)** Phenformin, **B)** IACS-010759 (IACS) **C)** Atorvastatin, and **D)** Fluvastatin. **E, F)** Dose response curves of DIPG-XIII GS and DGC treated with **E)** Phenformin, and **F)** Fluvastatin. **G-I)** Dose response curves of DIPG-XIII*P GS and DGC treated with **G)** Phenformin, **H)** Fluvastatin, and **I)** Atorvastatin. **J, K)** Dose response of PPK (murine H3K27M) GS and DGC treated with **J)** Phenformin, and **K)** Pitavastatin. **L)** Comparison of efficacy of various clinical statins (Atorvastatin, Fluvastatin, Lovastatin, Pitavastatin, and Simvastatin) in DIPG-007 GS. For all cell lines and treatment, cell viability was assayed using Cell Titer Glo 3 days post-treatment, except for DIPG-XIII*P in which cell viability was assessed 7 days post-treatment. All results are expressed as percent of control and mean ± SD or mean ± SEM.

**Supplementary Fig 7:**
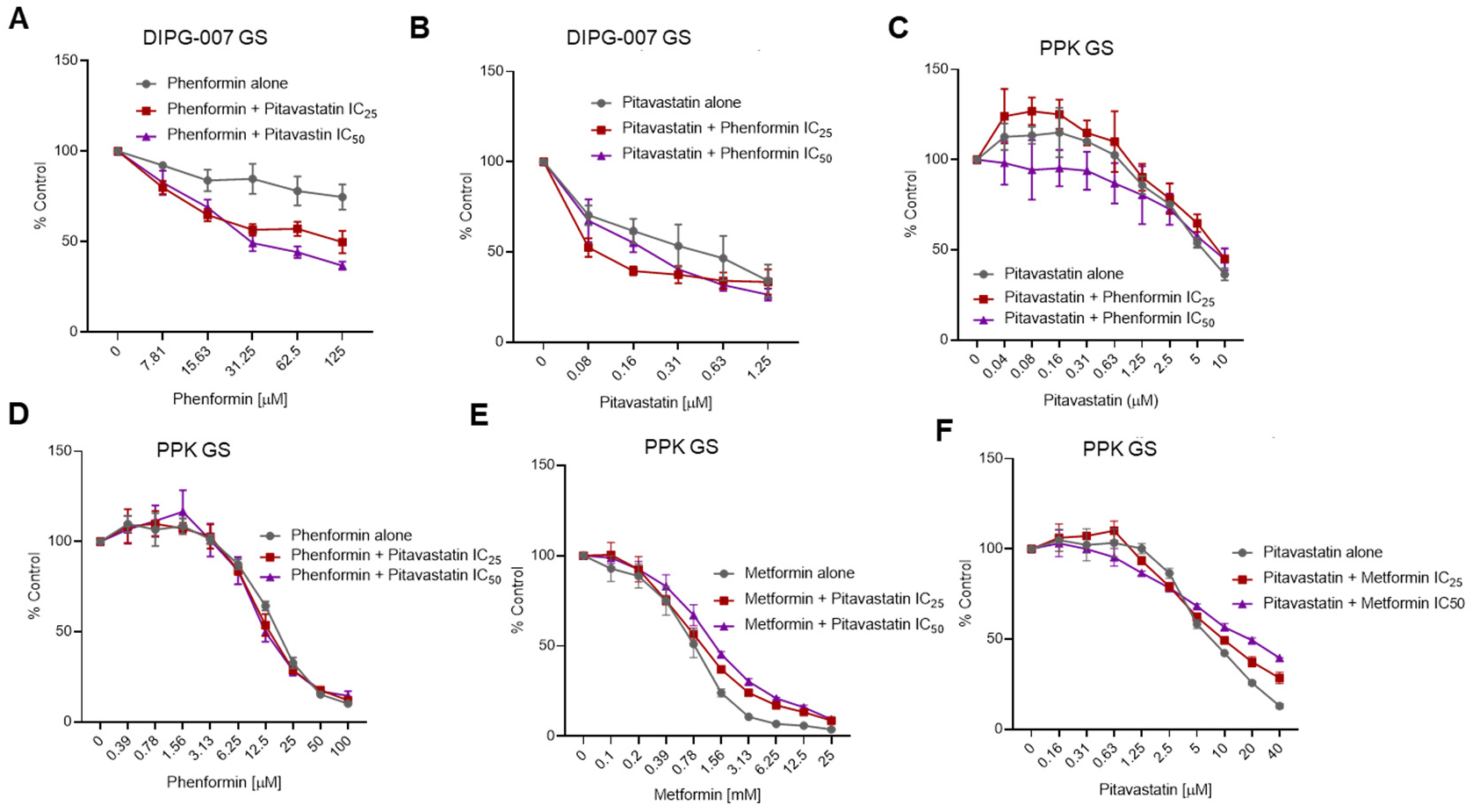
Combined metabolic targeting of cholesterol biosynthesis and OXPHOS in vitro. **A, B)** DIPG-007 GS treated for 3 days with different doses of Phenformin or Pitavastatin or in combination with the IC_25_, and IC_50_, of the respective combinatorial compound. **C-F)** Murine H3K27M PPK GS treated for 3 days with different doses of Phenformin, Metformin or Pitavastatin or in combination with the IC_25_, and IC_50_, of the respective combinatorial compound. Cell viability was assayed using Cell Titer Glo reagent and results expressed as a percentage of control (DMSO) and mean ± SD

**Supplementary Fig. 8:**
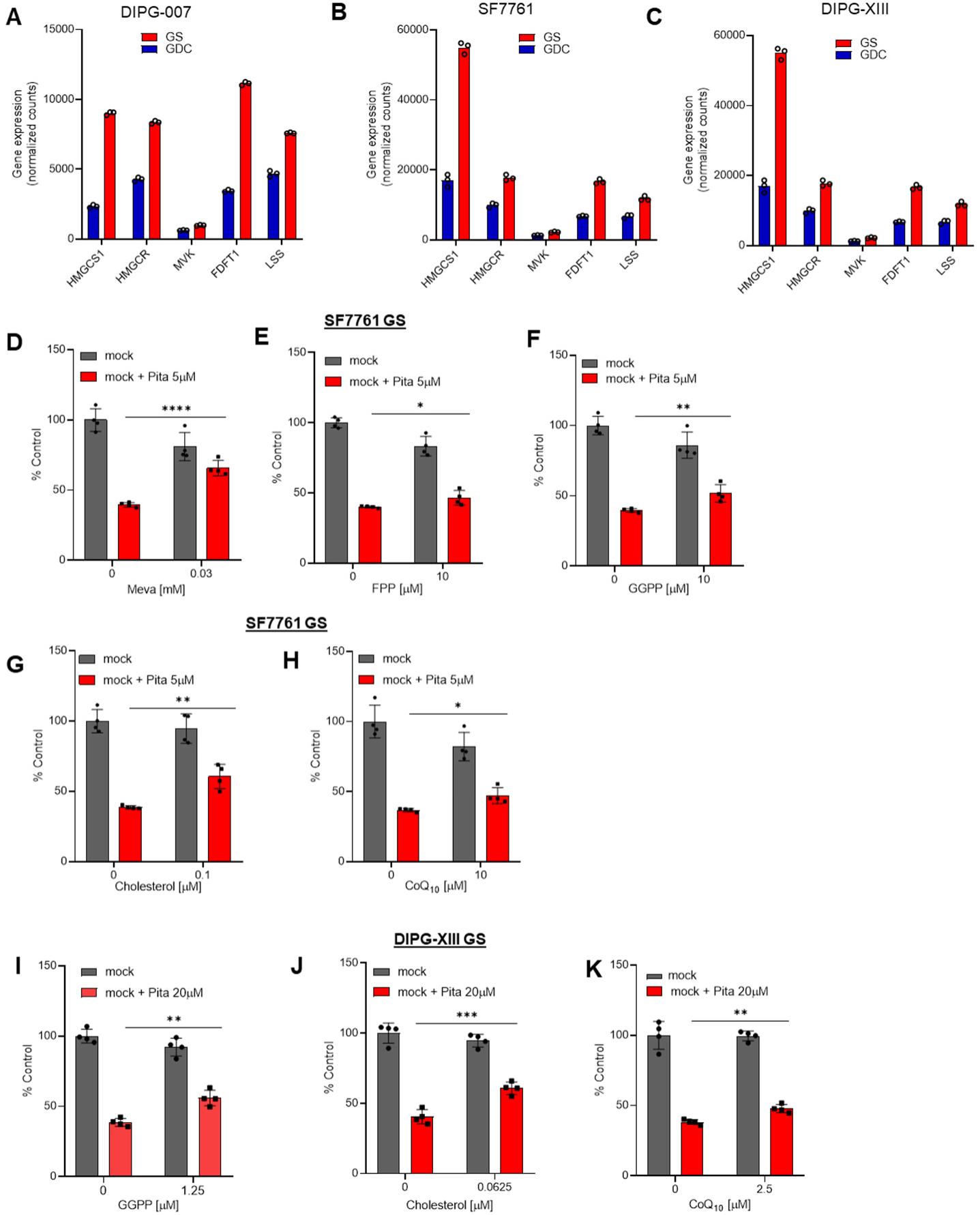
Dependency of DIPG GS on cholesterol biosynthesis. **A-C)** Relative expression of genes encoding enzymes involved in cholesterol biosynthesis in DIPG-007, SF7761, and DIPG-XIII GS and DGC counterparts. **D-H)** Cell viability of SF7761 GS following treatment with vehicle (0.4% DMSO) or Pitavastatin (Pita) with and without co-treated with Mevalonate (Meva), farnesyl pyrophosphate (FPP), geranylgeranyl pyrophosphate (GGPP), Cholesterol (Chol), and Coenzyme Q_10_ (CoQ_10_) at the indicated concentrations. Cell viability assayed at 3 days post-treatment using the Cell Titer Glo reagent and results expressed as a percent of control and mean ± SD. **I-K)** Cell viability of DIPG-XIII GS following treatment with vehicle (0.4% DMSO) or Pitavastatin (Pita) with and without co-treated with geranylgeranyl pyrophosphate (GGPP), Cholesterol (Chol), and Coenzyme Q1_0_ (CoQ_10_) at the indicated concentrations. Cell viability assayed at 3 days post-treatment using the Cell Titer Glo reagent and results expressed as a percent of control and mean ± SD (ns = not significant; * p < 0.05; ** p < 0.01; *** p < 0.001; **** p < 0.0001).

